# *Sox8* and *Sox9* act redundantly for ovarian-to-testicular fate reprogramming in the absence of *R-spondin1* in mouse sex reversals

**DOI:** 10.1101/2019.12.13.875443

**Authors:** Nainoa Richardson, Isabelle Gillot, Elodie P. Gregoire, Sameh A. Youssef, Dirk G. de Rooij, Alain de Bruin, Marie-Cécile De Cian, Marie-Christine Chaboissier

## Abstract

In mammals, testicular differentiation is initiated by transcription factors SRY and SOX9 in XY gonads, and ovarian differentiation involves R-spondin1 (RSPO1) mediated activation of WNT/β-catenin signaling in XX gonads. Accordingly, the absence of *RSPO1/Rspo1* in XX humans and mice leads to testicular differentiation and female-to-male sex reversal in a manner that does not require *Sry* or *Sox9* in mice. Here we show that an alternate testis-differentiating factor exists and that this factor is *Sox8*. Specifically, genetic ablation of *Sox8* and *Sox9* prevents ovarian-to-testicular reprogramming observed in XX *Rspo1* loss-of-function mice. Consequently, *Rspo1 Sox8 Sox9* triple mutant gonads developed as atrophied ovaries. Thus, SOX8 alone can compensate for the loss of SOX9 for Sertoli cell differentiation during female-to-male sex reversal.

## Introduction

During primary sex determination in mammals, a common precursor organ, the bipotential gonad, develops as a testis or ovary. In humans and mice, testicular development begins when SRY and SOX9 are expressed in the bipotential XY gonad. These transcription factors promote supporting cell progenitors to differentiate as Sertoli cells and form sex cords (Chaboissier, Kobayashi et al. 2004, Barrionuevo, Bagheri-Fam et al. 2006, Gonen, Futtner et al. 2018), and this triggers a cascade of signaling events that are required for the differentiation of other cell populations in the testis (Koopman, Gubbay et al. 1991, Vidal, Chaboissier et al. 2001). In XX embryos, the bipotential gonad differentiates as an ovary through a process that requires RSPO1-mediated activation of canonical WNT/β-catenin (CTNNB1) signaling in somatic cells (Parma, Radi et al. 2006, Chassot, Ranc et al. 2008). Ovarian fate also involves activation of FOXL2, a transcription factor that is required in post-natal granulosa cells (Schmidt, Ovitt et al. 2004, Ottolenghi, Omari et al. 2005), which organize as follicles during embryogenesis in humans and after birth in mice (McGee and Hsueh 2000, Mork, Maatouk et al. 2012). For complete differentiation of testes or ovaries, an active repression of the opposite fate is necessary (Kim, Kobayashi et al. 2006). Inappropriate regulation within the molecular pathways governing sex determination can lead to partial or complete sex reversal phenotypes and infertility (Wilhelm, Washburn et al. 2009).

Studies in humans and mice have shown that the pathway initiated by SRY/SOX9 or RSPO1/WNT/β-catenin signaling are indispensable for sex specific differentiation of the gonads. For example, in XY humans, *SRY* or *SOX9* loss-of-function mutations prevent testis development (Houston, Opitz et al. 1983, Berta, Hawkins et al. 1990). In mice, XY gonads developing without SRY or SOX9 lack Sertoli cells and seminiferous tubules and differentiate as ovaries that contain follicles (Lovell-Badge and Robertson 1990, Chaboissier, Kobayashi et al. 2004, Barrionuevo, Bagheri-Fam et al. 2006, Lavery, Lardenois et al. 2011, Kato, Miyata et al. 2013), indicating *Sry*/*Sox9* requirement. In XX humans and mice, *SRY*/*Sry* or *SOX9/Sox9* gain-of-function mutations promote Sertoli cell differentiation and testicular development (Sinclair, Berta et al. 1990, Koopman, Gubbay et al. 1991, Huang, Wang et al. 1999, Bishop, Whitworth et al. 2000, Vidal, Chaboissier et al. 2001), indicating that SRY/SOX9 function is also sufficient for male gonad differentiation.

With respect to the ovarian pathway, homozygous loss-of-function mutations for *RSPO1/Rspo1* trigger female-to-male sex reversal in XX humans and mice (Parma, Radi et al. 2006, Chassot, Ranc et al. 2008). In XX *Rspo1* or *Wnt4* mutant mice, Sertoli cells arise from a population of embryonic granulosa cells (pre-granulosa cells) that precociously exit their quiescent state, differentiate as mature granulosa cells, and reprogram as Sertoli cells (Chassot, Ranc et al. 2008, Maatouk, Mork et al. 2013). The resulting gonad is an ovotestis containing seminiferous tubule-like structures with Sertoli cells and ovarian follicles with granulosa cells, indicating that SRY is dispensable for testicular differentiation. In addition, stabilization of WNT/CTNNB1 signaling in XY gonads leads to male-to-female sex reversal (Maatouk, DiNapoli et al. 2008, Harris, Siggers et al. 2018). Thus, RSPO1/WNT/CTNNB1 signaling is required for ovarian differentiation and female development in humans and mice.

Given the prominent role of SOX9 for testicular development (Chaboissier, Kobayashi et al. 2004, Barrionuevo, Georg et al. 2009), it was hypothesized that SOX9 is responsible for Sertoli cell differentiation in XX gonads developing without *RSPO1*/*Rspo1*. This hypothesis was tested by co-inactivation of *Rspo1* or *Ctnnb1* and *Sox9* in *Rspo1 Sox9* (Lavery, Chassot et al. 2012) and in *Ctnnb1 Sox9* double mutant mice (Nicol and Yao 2015). Unexpectedly, XY and XX *Rspo1* or *Ctnnb1* mutant gonads lacking *Sox9* exhibited Sertoli cells organized as testis cords (Lavery, Chassot et al. 2012, Nicol and Yao 2015). Specifically, gonads in XX *Rspo1 Sox9* double mutant mice developed as ovotestes as in XX *Rspo1* single mutants, and XY *Rspo1 Sox9* mutant mice developed hypoplastic testes capable of supporting the initial stages of spermatogenesis. These outcomes indicate that at least one alternate factor can promote testicular differentiation in *Rspo1* mutant mice also lacking *Sox9* in XY mice, and lacking both *Sry* and *Sox9* in XX animals. This or these factors remained to be identified.

Among the candidate genes that could promote testicular differentiation in the absence of *Sry* and *Sox9* are the other members of the *SoxE* group of transcription factors that includes *Sox9*: *Sox8* and *Sox10* (Lavery, Chassot et al. 2012, Nicol and Yao 2015). However, *Sox10* expression in testes depends on *Sox8* and *Sox9* (Georg, Barrionuevo et al. 2012), and *Sox10* loss-of-function mice are fertile (Peirano and Wegner 2000, Britsch, Goerich et al. 2001), suggesting that *Sox10* would not be the best candidate gene. For *Sox8*, loss-of-function analyses in XY gonads show testicular development, indicating that *Sox8* is not required for Sertoli cell differentiation during embryonic development (Sock, Schmidt et al. 2001). However, a *Sox8*-null background enhanced the penetrance of the testis-to-ovary sex reversal phenotype in mice with reduced *Sox9* expression (Chaboissier, Kobayashi et al. 2004), suggesting that *Sox8* supports the function of *Sox9*.

Furthermore, in XY *Sox9* single mutant and XY *Sox8 Sox9* double mutant mice where *Sox9* is inactivated after sex determination, the single and double mutant mice form testis cords containing Sertoli cells, these cells loose their identity and begin to express granulosa cell markers like FOXL2 (Barrionuevo, Georg et al. 2009, Georg, Barrionuevo et al. 2012, Barrionuevo, Hurtado et al. 2016). In addition, the double *Sox8,9*-null Sertoli cells become apoptotic, leading to a complete degeneration of the seminiferous tubules. This indicated that a concerted effort by *Sox8* and *Sox9* is required in XY gonads for the maintenance of Sertoli cells after sex determination.

Although *Sox8* expression is dispensable for Sertoli cell differentiation in XY gonads, it may have a key role for testicular differentiation in XX sex reversal gonads or in cases of *Sox9*-independent testicular differentiation in XY gonads. This led us to hypothesize that *Sox8* can compensate for loss of *Sox9* and is the alternate factor capable of: (*i*) triggering sex reversal in XX *Rspo1* knockout gonads lacking *Sry* and *Sox9*, and (*ii*) promoting testicular development in XY *Rspo1* knockout gonads lacking *Sox9*.

To test this hypothesis, we have generated triple *Rspo1, Sox8,* and *Sox9* loss-of-function mutant mice models. We show here that *Sox8* and *Sox9* are individually dispensable for testicular development in XY and XX mice lacking *Rspo1*, indicating the presence of redundant testicular pathways. In the absence of both *Sox* factors, Sertoli cell differentiation is precluded and XY and XX *Rspo1 Sox8 Sox9* triple mutants develop atrophied ovaries. Together, our data show that *Sox8* or *Sox9* is required to induce testicular development in XY and XX mice lacking *Rspo1*.

## Results

### *Rspo1*, *Sox8,* and *Sox9* are expressed independently

We first performed expression analyses for *Rspo1*, *Sox8*, and *Sox9* in control and mutant gonads. We chose to study embryonic day 17.5 (E17.5) fetal gonads, when testis cords form in *Rspo1* sex reversal mice, and juvenile postnatal day 10 (P10) gonads, when gonadal fate is likely to be set (Lavery, Chassot et al. 2012). In XY gonads, *Rspo1* is mostly localized to the coelomic epithelium at E17.5 and to the tunica albuginea at P10 (*Figure 1A,C*). In fetal ovaries, *Rspo1* is expressed in the somatic cells at E17.5 (*Figure 1B*) and down-regulated after birth, as shown in post-natal P10 ovaries (*Figure 1D*). In XY and XX mice lacking *Sox8* and *Sox9* (i.e., *Sox8^−/−^ Sox9^fl/fl^; Sf1:cre^Tg/+^*, referred to as *Sox8^KO^ Sox9^cKO^* double mutants), high *Rspo1* expression levels were observed in embryonic gonads, indicating ovarian differentiation (*Figure 1E,F*). This was later confirmed by immunostainings for FOXL2, a granulosa cell marker (*Figure 3G and Figure 4E*). These data confirmed that although *Rspo1* is expressed in both XY and XX gonads, robust *Rspo1* expression in cells throughout the gonad is a feature of ovarian development in fetuses.

**Figure 1.**
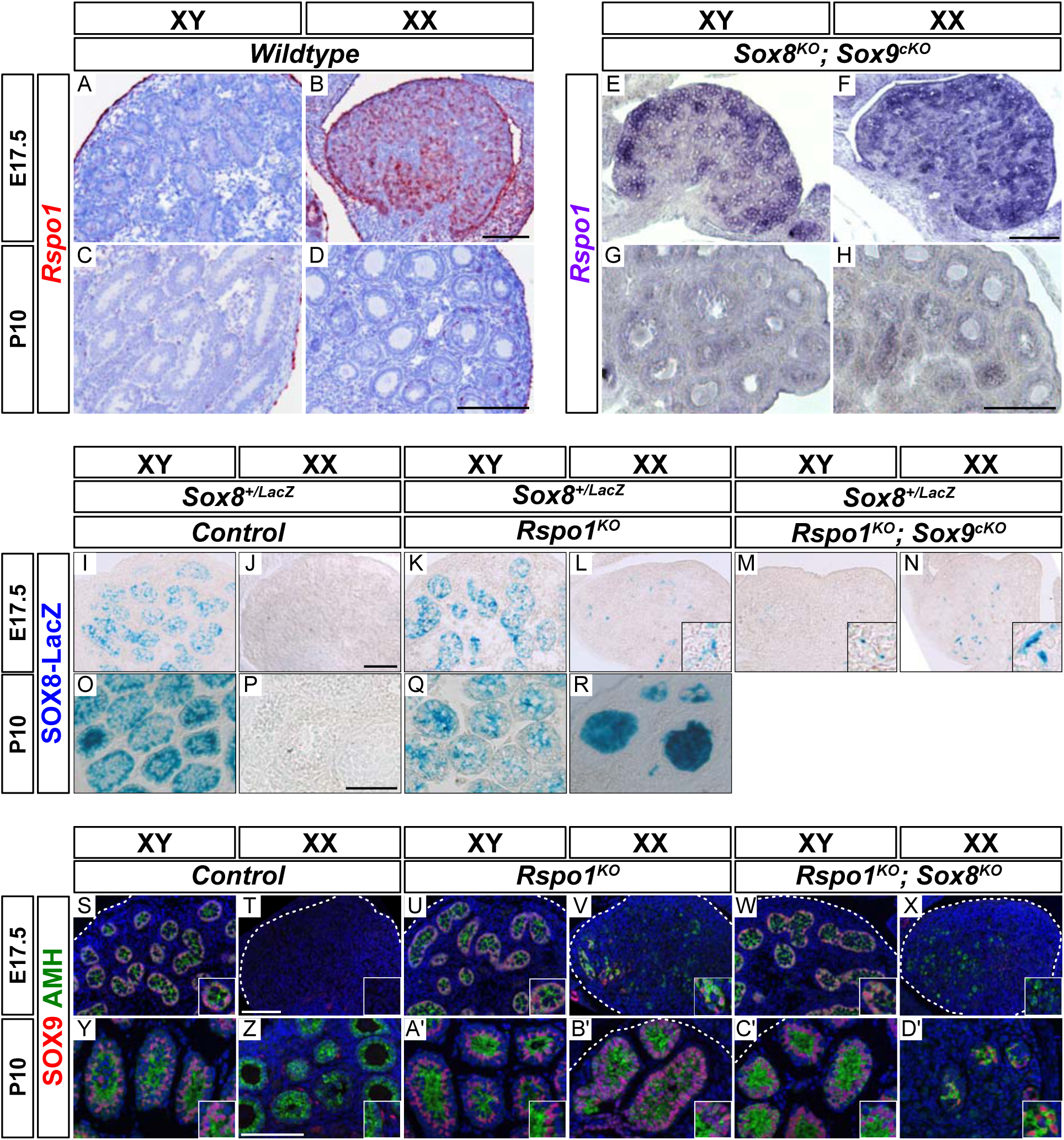
Expression of *Rspo1*, *Sox8*, and *Sox9* in E17.5 and P10 gonads. Expression of *Rspo1*, as revealed by *in situ* hybridizations (A-H), SOX8-LacZ expression as visualized by β-galactosidase staining (I-R), and SOX9 expression as revealed by immunostainings (S-D’) on gonadal sections from embryonic day 17.5 (E17.5) and 10 days post-natal (P10) mice. In XY wild-type testes, *Rspo1* is mainly expressed in the coelomic epithelium (A) and tunica albuginea (C). In XX wildtype ovaries, *Rspo1* is expressed throughout the gonad at E17.5 (B), and down-regulated in post-natal animals, as shown at P10 (D). In XY and XX *Sox8^−/−^*; *Sox9^flox/flox^*; *Sf1:cre^Tg/+^* (*Sox8^KO^ Sox9^cKO^*) mutant mice, the *Rspo1* expression profile (E-H) is similar to wildtype ovaries (B, D), suggesting an ovarian fate. This was later confirmed by immunostaining experiments for FOXL2, a granulosa cell maker (Figures 3G and 4E). To visualize the expression of SOX8-LacZ, control and mutant mice contained a *Sox8^+/−^* (*Sox8^+/LacZ^*) heterozygous background (I-R). In XY control testes (I, O) and in XY *Rspo1^KO^* gonads developing as testes (K, Q), SOX8-LacZ expression was found in testis cords at E17.5 and P10. In XX mice, SOX8-LacZ expression is absent in XX control ovaries (J, P), but found in XX *Rspo1^KO^* female-to-male sex reversal gonads (L, R). SOX8-LacZ is also expressed in the absence of *Sox9* in XY and XX *Rspo1^KO^ Sox9^cKO^* gonads at E17.5 (M, N), which is in agreement with our previous study in P12 mice (see (Lavery, Chassot et al. 2012). SOX9 expression was found in XY control testes (S, Y), and in XY *Rspo1^KO^* gonads developing as testes (U, A’). Co-immunolabeling with AMH confirmed the identity of Sertoli cells. As shown, SOX9-, AMH-positive testis cords are found in XX *Rspo1^KO^* sex reversal gonads at E17.5 and P10 (V, B’). SOX9 is also expressed in absence of *Sox8* in XY *Rspo1−/−*; *Sox8*^*−/−*^ (*Rspo1^KO^ Sox8^KO^*) gonads developing as testes at E17.5 and P10 (W, C’), and in XX *Rspo1^KO^ Sox8^KO^* gonads developing as ovotestes at P10 (D’). In XX control mice, SOX9 and AMH expression is absent in fetal ovaries (T). In post-natal female animals, SOX9 is expressed by theca cells, which are AMH-negative (Z). All scale bars 100μm.

Next, we carried out *Sox8* expression analyses in *Rspo1^−/−^* mutants (referred to as *Rspo1^KO^* mutants) at E17.5 and P10, and in XY and XX *Rspo1^−/−^*; *Sox9^fl/fl^; Sf1:cre^Tg/+^* gonads (referred to as *Rspo1^KO^ Sox9^cKO^* double mutants) at E17.5, given that *Sox8* expression was previously shown in XY and XX *Rspo1^KO^ Sox9^cKO^* double mutants around P10 (Lavery et al., 2012). Here, we took advantage of the *LacZ* reporter inserted into the *Sox8* mutant allele (Sock, Schmidt et al. 2001). This approach was possible in *Rspo1* mutant mice also carrying a *LacZ* reporter (Chassot, Ranc et al. 2008), since β-galactosidase activity is not detectable in *Rspo1* heterozygous and homozygous gonads at E13.5 (*Figure 1–figure supplement 1B,C*), a time when *Rspo1* is highly expressed (Parma, Radi et al. 2006). The lack of intracellular β-galactosidase is due to the retention of the secretion signal in RSPO1-β-galactosidase protein expressed by the *Rspo1* knock-out allele (Chassot, Ranc et al. 2008).

We observed SOX8-LacZ expression in XX *Rspo1^KO^* and XY and XX *Rspo1^KO^ Sox9^cKO^* gonads in fetuses containing a *Sox8^+/LacZ^* background at E17.5 (*Figure 1L-N*). At this stage, LacZ-positive cells were not abundant, highlighting the beginning of Sertoli cell differentiation and sex cord formation in ovotestes as described in (Chassot, Ranc et al. 2008, Maatouk, Mork et al. 2013). In testis cords of XY control and XY *Rspo1^KO^* mice at E17.5 and P10, SOX8-LacZ expression was readily detectable (*Figure 1I,O,K,Q*), as was also the case in XX *Rspo1^KO^* sex reversal gonads at P10 (*Figure 1R*). In contrast, control ovaries lacked SOX8-LacZ expression (*Figure 1J,P*). Thus, these data corroborate our previous study showing that *Sox8* is expressed in the absence of *Rspo1* and *Sox9* in XY mice, and additionally in absence of *Sry* in XX animals during juvenile stages (Lavery, Chassot et al. 2012).

Next, immunostainings revealed SOX9-positive testis cords in XY *Rspo1^KO^* testes (*Figure 1U,A’*), XX *Rspo1^KO^* ovotestes (*Figure 1V,B’*), as in control testes (*Figure 1S,Y*), as previously described (Chassot, Ranc et al. 2008). Co-immunolabeling with AMH confirmed the identity of Sertoli cells, since AMH-positive granulosa cells do not express *Sox9*, and given that ovarian steroidogenic theca cells expressing *Sox9* are AMH-negative (*Figure 1Z*). In addition, deletion of *Sox8* did not alter the expression of *Sox9* in XY or XX *Rspo1^KO^* gonads (i.e., in *Rspo1^KO^ Sox8^KO^* gonads) (*Figure 1W,C’,X,D’*). Altogether, our results show that *Sox8* and *Sox9* are expressed in the absence of each other in *Rspo1* mutant gonads during testis cord development.

### Ablation of *Rspo1* and *Sox8* does not impair testis differentiation

Next, we asked whether inactivation of both *Rspo1* and *Sox8* would impact gonad development in XY *Rspo1^KO^ Sox8^KO^* double mutants. In XY *Rspo1^KO^ Sox8^KO^* mice, the anogenital distance in adult P40 animals was comparable to XY control males (*Figure 2–figure supplement 1A,B*). In contrast, XX control females exhibited a short anogenital distance (*Figure 2–figure supplement 1E*). Internally, XY *Rspo1^KO^ Sox8^KO^* mice developed epididymides, vasa deferensia, seminal vesicles and prostate, as in control males (*Figure 2–figure supplement 1F,G*). Histological analyses by PAS staining revealed seminiferous tubules with no obvious defects in P10 and P40 XY *Rspo1^KO^ Sox8^KO^* animals (*Figure 2–figure supplement 1L and Figure 2G*), and these mice were fertile. Testicular development in XY *Rspo1^KO^ Sox8^KO^* mice was confirmed by immunostaining experiments on embryonic (E17.5) and post-natal (P10, and P40) gonads that contained SOX9-and DMRT1-positive Sertoli cells forming testicular sex cords and seminiferous tubules (*Figure 1W,C’, Figure 2–figure supplement 1Q,V, and Figure 2L,Q*). DMRT1 expression was also observed in germ cells, which are TRA98-positive (*Figure 2L,Q, and Figure 2–figure supplement 1Q,V*) (Matson, Murphy et al. 2010). In contrast with XY *Rspo1^KO^ Sox8^KO^* mice, gonads in XX control mice exhibited follicles containing granulosa cells expressing FOXL2 (*Figure 2–figure supplement 1O,T,Y and Figure 2J,O,T*). Thus, loss of both *Rspo1* and *Sox8* does not impair testis differentiation.

**Figure 2.**
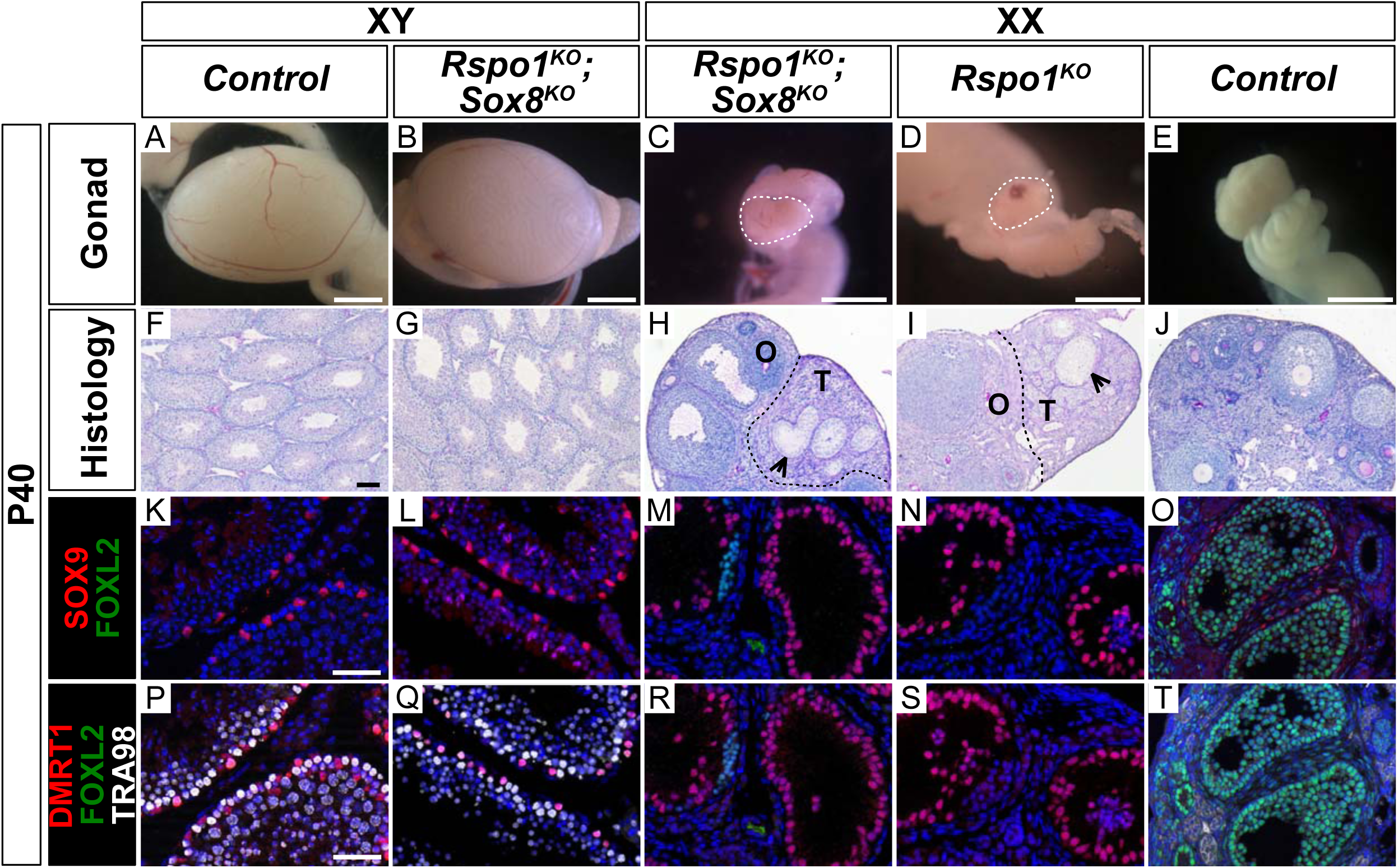
Testis and ovotestis development in adult XY and XX *Rspo1^KO^ Sox8^KO^* double mutant mice. Macroscopic view of gonads from adult P40 mice (A-E) (Scale bars 1.5mm), histology as revealed by PAS staining on gonadal sections (F-J) (Scale bars 100μm), and immunostainings of SOX9 (Sertoli cell marker, in red) (K-O), FOXL2 (granulosa cell marker, in green) (K-T), DMRT1 (Sertoli and germ cell marker, in red) (P-T), TRA98 (germ cell marker, in white) (P-^), and DAPI (nuclear marker, in blue) (K-T) on gonadal sections (Scale bars 50μm). Inactivation of both *Rspo1* and *Sox8* in XY *Rspo1^KO^ Sox8^KO^* double mutant mice did not cause a sex reversal (A, B). XY *Rspo1^KO^ Sox8^KO^* gonads developed as testes with seminiferous tubules (G) containing SOX9-and DMRT1-postive Sertoli cells (L, Q), as in control testes (F, K, P). As shown, XX control ovaries developed follicles (J) containing FOXL2-positive granulosa cells (O, T). Adult ovotestes in XX *Rspo1^KO^ Sox8^KO^* mice (C, H) were indistinguishable from XX *Rspo1^KO^* mice (D, I). These gonads contained an ovarian “O” compartment with follicles and a testicular “T” compartment with seminiferous tubule-like structures, as indicated by arrowheads (H, I). The seminiferous tubule-like structures in XX *Rspo1^KO^ Sox8^KO^* and XX *Rspo1^KO^* ovotestes contained SOX9-and DMRT1-positive Sertoli cells (M, N, R, S), as in control testes (K, P), but lacked TRA98-postive germ cells (R, S).

We then asked if gonads in XX *Rspo1^KO^ Sox8^KO^* mice developed as ovaries or as ovotestes, as in XX *Rspo1^KO^* and XX *Rspo1^KO^ Sox9^cKO^* mice (Lavery, Chassot et al. 2012). In XX *Rspo1^KO^ Sox8^KO^* and XX *Rspo1^KO^* gonads at P10 respectively, testis cords and seminiferous tubules devoid of germ cells were apparent (*Figure 2–figure supplement 1R,W,S,X*). This suggested a delay in ovo-testicular development in double mutant gonads. Indeed, by P40, both XX *Rspo1^KO^* and XX *Rspo1^KO^ Sox8^KO^* mice were essentially indistinguishable with respect to gonad morphology (*Figure 2C,D*), reproductive tract development (*Figure 2–figure supplement 1H,I*), ovo-testicular organization (*Figure 2H,I*), and the presence of SOX9-and DMRT1-positive Sertoli cells in the testicular area (*Figure 2M,N,R,S*). Altogether, studies performed in *Rspo1^KO^ Sox8^KO^* mice demonstrate that like *Sox9* (Lavery, Chassot et al. 2012), *Sox8* is dispensable for testicular development in XY and XX *Rspo1^KO^* gonads. Moreover, our data suggests that SOX9 likely compensates for the loss of *Sox8* in *Rspo1^KO^ Sox8^KO^* double mutants.

### Ovarian precocious differentiation occurs in XX and XY *Rspo1^KO^ Sox8^KO^ Sox9c^KO^* fetuses

Our genetic mouse models allowed us to investigate gonadal fate in XY and XX *Rspo1^KO^* mice lacking both *Sox8* and *Sox9* (i.e., in XY and XX *Rspo1^KO^ Sox8^KO^ Sox9^cKO^* triple mutant mice). We first studied gonads in E17.5 fetuses, which is when differentiated granulosa cells reprogram as Sertoli cells in XX *Rspo1^KO^* gonads (Maatouk, Mork et al. 2013). As shown, XX control gonads contained granulosa cells expressing FOXL2, but not Sertoli cells expressing SOX9 or DMRT1 (*Figure 3–figure supplement 1A and Figure 3F*), indicating ovarian development. The granulosa cells remained quiescent, as evidenced by expression of the mitotic arrest marker CDKN1B (also known as P27) throughout the E17.5 gonad, and the absence of AMH expression indicated that these cells were fetal or pre-granulosa cells (*Figure 3A*) (Maatouk, Mork et al. 2013).

**Figure 3.**
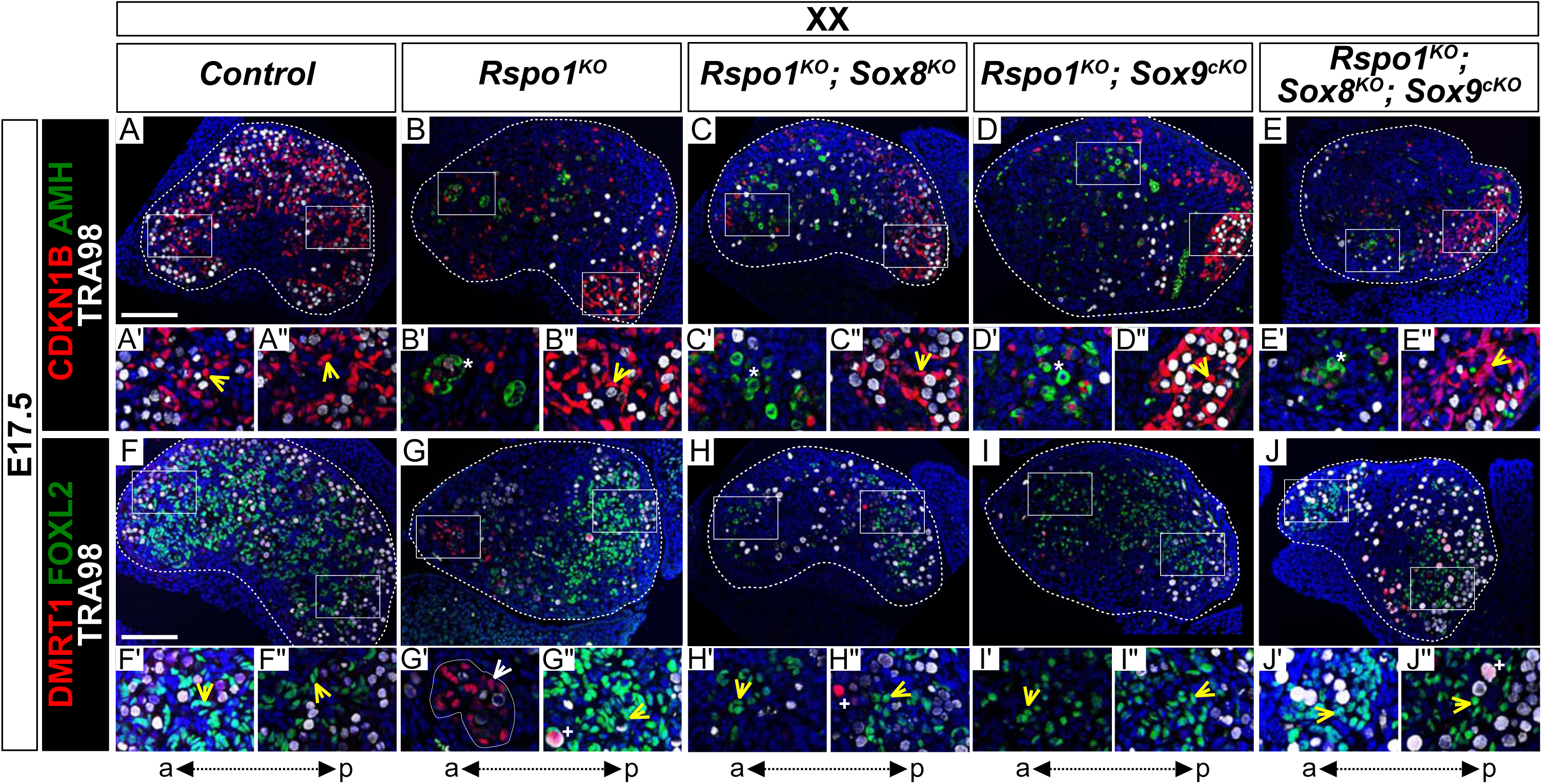
Precocious granulosa cell differentiation in XX *Rspo1^KO^ Sox8^KO^ Sox9^cKO^* triple mutant fetuses at E17.5. Immunofluorescence of CDKN1B (P27) (mitotic arrest marker, in red) (A-E), AMH (Sertoli marker and mature granulosa cell marker, in green) (A-E), DMRT1 (Sertoli and germ cell marker, in red) (F-J), FOXL2 (granulosa cell marker, in green) (F-J), TRA98 (A-J) (germ cell marker, in white) (A-J), and DAPI (nuclear marker, in blue) (A-J) on gonadal sections from E17.5 fetuses (main panel scale bar 100μm). The anterior “a” and posterior “p” axis is shown below each column. For main panels A-J, highlighted anterior and posterior areas are shown in the respective single and double primed letter panels. Yellow arrowheads indicate granulosa cells expressing CDKN1B or FOXL2, asterisks indicate cells expressing AMH, white arrowheads indicate Sertoli cells expressing DMRT1, and plus symbols indicate germ cells expressing DMRT1 and TRA98. Gonads in XX control fetuses developed as ovaries, as shown by FOXL2 and CDKN1B expression in pre-granulosa cells (A’ and F’, yellow arrowheads). These fetal ovaries lacked mature granulosa cells expressing AMH (A). In contrast, XX *Rspo1^−/−^* (*Rspo1^KO^*), XX *Rspo1^−/−^*; *Sox8^−/−^* (*Rspo1^KO^ Sox8^KO^*), XX *Rspo1^−/−^*; *Sox9^flox/flox^*; *Sf1:cre^Tg/+^* (*Rspo1^KO^ Sox9^cKO^*), and XX *Rspo1^KO^ Sox8^KO^ Sox9^cKO^* gonads exhibited down-regulation of CDKN1B (B-E) and ectopic AMH expression in the anterior area (B’-E’, asterisks), indicating Sertoli cells or mature granulosa cells. However, while XX *Rspo1^KO^* gonads contained Sertoli cells expressing DMRT1 (G’, white arrowheads), these cells were rare in XX double and triple mutants (H-J) (1 out of 8 XX triple mutant gonads studied from n=4 fetuses). Note that some DMRT1-positive cells are germ cells expressing TRA98 (G”, H”, J”, plus symbols). Thus, while granulosa cells differentiate precociously in XX *Rspo1^KO^* gonads lacking *Sox8* and/or *Sox9* at E17.5, these cells have not yet reprogrammed as Sertoli cells in XX *Rspo1^KO^ Sox8^KO^* and XX *Rspo1^KO^ Sox9^cKO^* mice. In XX triple mutant fetuses, granulosa cell reprogramming as Sertoli cells may be delayed, or blocked.

In contrast, CDKN1B is down-regulated in the anterior area of XX *Rspo1^KO^ Sox8^KO^ Sox9^cKO^* triple mutant gonads (*Figure 3E*), as in XX *Rspo1^KO^* single, as well as in XX *Rspo1^KO^ Sox8^KO^* and XX *Rspo1^KO^ Sox9^cKO^* double mutants (*Figure 3B,C,D*) (Maatouk, Mork et al. 2013). In addition, these mutants contained cells expressing AMH (*Figure 3B’,C’,D’,E’ asterisks*), indicating precocious granulosa cell differentiation, as previously described (Maatouk, Mork et al. 2013). However, while SOX9-and DMRT1-positive, TRA98-negative Sertoli cells were readily detectable in the anterior area of the XX *Rspo1^KO^* gonads (*Figure 3–figure supplement 1B’ and Figure 3G’ white arrowheads*), these cells were noticeably absent or rare in XX *Rspo1^KO^ Sox8^KO^ Sox9^cKO^* triple mutant fetuses (1 out of 8 XX triple mutant gonads studied from n=4 fetuses) (*Figure 3–figure supplement 1E and Figure 3J*). This was also the case in XX *Rspo1^KO^ Sox8^KO^* and XX *Rspo1^KO^ Sox9^cKO^* double mutants (*Figure 3–figure supplement 1 C,D and Figure 3H,I*). Together with these observations, quantification of immunostained cells expressing DMRT1, FOXL2, and CDKN1B per gonadal section area demarcated by DAPI confirmed the lack of Sertoli cells and presence of granulosa cells in XX double and triple mutant gonads at E17.5 (*Figure 4A,C,E*).

**Figure 4.**
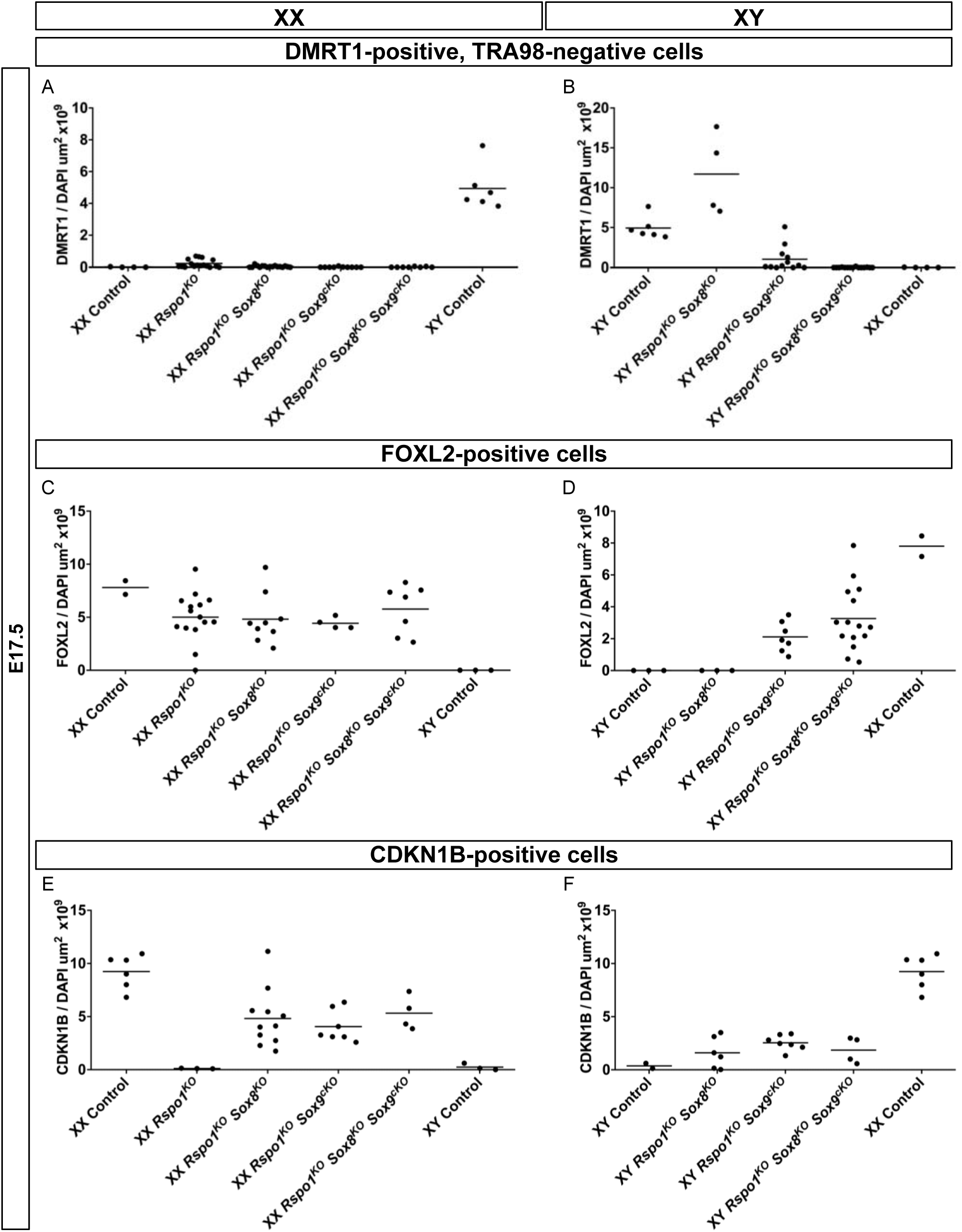
Quantification of immunostained cells expressing DMRT1, FOXL2, and CDKN1B at E17.5. Histogram showing the distribution of immunostained Sertoli expressing DMRT1 and negative for TRA98 (A-B), granulosa cells expressing FOXL2 (C-D), and non-dividing cells expressing CDKN1B (E-F) per gonad section area (μm^2^), as demarcated by DAPI, and as shown in Figures 3 and 4. As shown, XX control fetuses contained granulosa cells expressing FOXL2 (C), and lacked cells Sertoli cells expressing DMRT1 (A). In contrast, XX *Rspo1^−/−^* (*Rspo1^KO^*) gonads have started to develop DMRT1-positive Sertoli cells (A), while still maintaining a partial FOXL2-positive ovarian identity (C). In the XX single mutant, CDKN1B is down-regulated compared with XX control fetal ovaries, indicating that some pre-granulosa cells have precociously existed their mitotic arrest. Similarly, XX *Rspo1^−/−^*; *Sox8^−/−^* (*Rspo1^KO^ Sox8^KO^*), XX *Rspo1^−/−^*; *Sox9^flox/flox^*; *Sf1:cre^Tg/+^* (*Rspo1^KO^ Sox9^cKO^*), and XX *Rspo1^KO^ Sox8^KO^ Sox9^cKO^* gonads exhibited granulosa cells (C, E), but these mutant gonads lacked Sertoli cells expressing DMRT1 (A). Gonads from XY *Rspo1^KO^ Sox8^KO^* fetuses contained DMRT1-positive Sertoli cells and lacked granulosa cells expressing FOXL2, as in control XY fetuses (B, D). Gonads in XY *Rspo1^KO^ Sox9^cKO^* fetuses also exhibited DMRT1-positive Sertoli cells, which were rare in XY *Rspo1^KO^ Sox8^KO^ Sox9^cKO^* gonads (B). Gonads from both XY *Rspo1^KO^ Sox9^cKO^* and XY triple mutant fetuses contained FOXL2-and CDKN1B-positive cells, indicating granulosa cells.

In addition to the presence of mature granulosa cells, gonads in the XX single, double, and triple mutant fetuses also exhibited NR5A1-and HSD3β-positive cells (*Figure 3–figure supplement 1G,H,I,J*), which were absent in XX control ovaries (*Figure 3–figure supplement 1F*) (Chassot, Ranc et al. 2008, Lavery, Chassot et al. 2012). Thus, these data indicated that ablation of *Sox8* and/or *Sox9* in XX fetuses lacking *Rspo1* does not prevent the appearance of steroidogenic cells and precocious granulosa differentiation, two characteristics of XX *Rspo1^KO^* gonads (Chassot, Ranc et al. 2008, Maatouk, Mork et al. 2013).

We then examined the phenotype of *Rspo1^KO^ Sox8^KO^ Sox9^cKO^* gonads in E17.5 XY fetuses. As shown, XY *Rspo1^KO^ Sox8^KO^* double mutant gonads contained SOX9-and DMRT1-positive Sertoli cells forming testis cords, as in control fetal testes (*Figure 5–figure supplement 1A,B and Figure 5E,F*). Also, XY *Rspo1^KO^ Sox9^cKO^* gonads exhibited DMRT1-positive testis chords (*Figure 5G,G’*), which were more pronounced than testis cords in XX *Rspo1^KO^* and XX *Rspo1^KO^ Sox9^cKO^* gonads at this stage (*Figure 3G,G’,I*) (Lavery, Chassot et al. 2012). Thus, in XY fetuses lacking *Rspo1*, inactivation of one *Sox* gene is dispensable for Sertoli cells. However, in fetuses lacking both *Sox8* and *Sox9* in XY triple mutant gonads, Sertoli cells expressing SOX9 or DMRT1 were not readily obvious (6 of 6 XY triple mutant gonads studied from n=3 fetuses) (*Figure 5–figure supplement 1D and Figure 5H*). Instead, XY triple mutant gonads exhibited FOXL2-positive pre-granulosa cells (*Figure 5H,H’,H’’’, yellow arrowheads*), and AMH expression suggested that some mature granulosa cells were present (*Figure 5D’ asterisk*). Quantification of cells expressing expressing DMRT1, FOXL2, and CDKN1B confirmed these observations (*Figure 4B,D,F*). Like XX triple mutants, XY triple mutants also contained steroidogenic cells expressing NR5A1 and HSD3β (*Figure 5–figure supplement 1H*).

**Figure 5.**
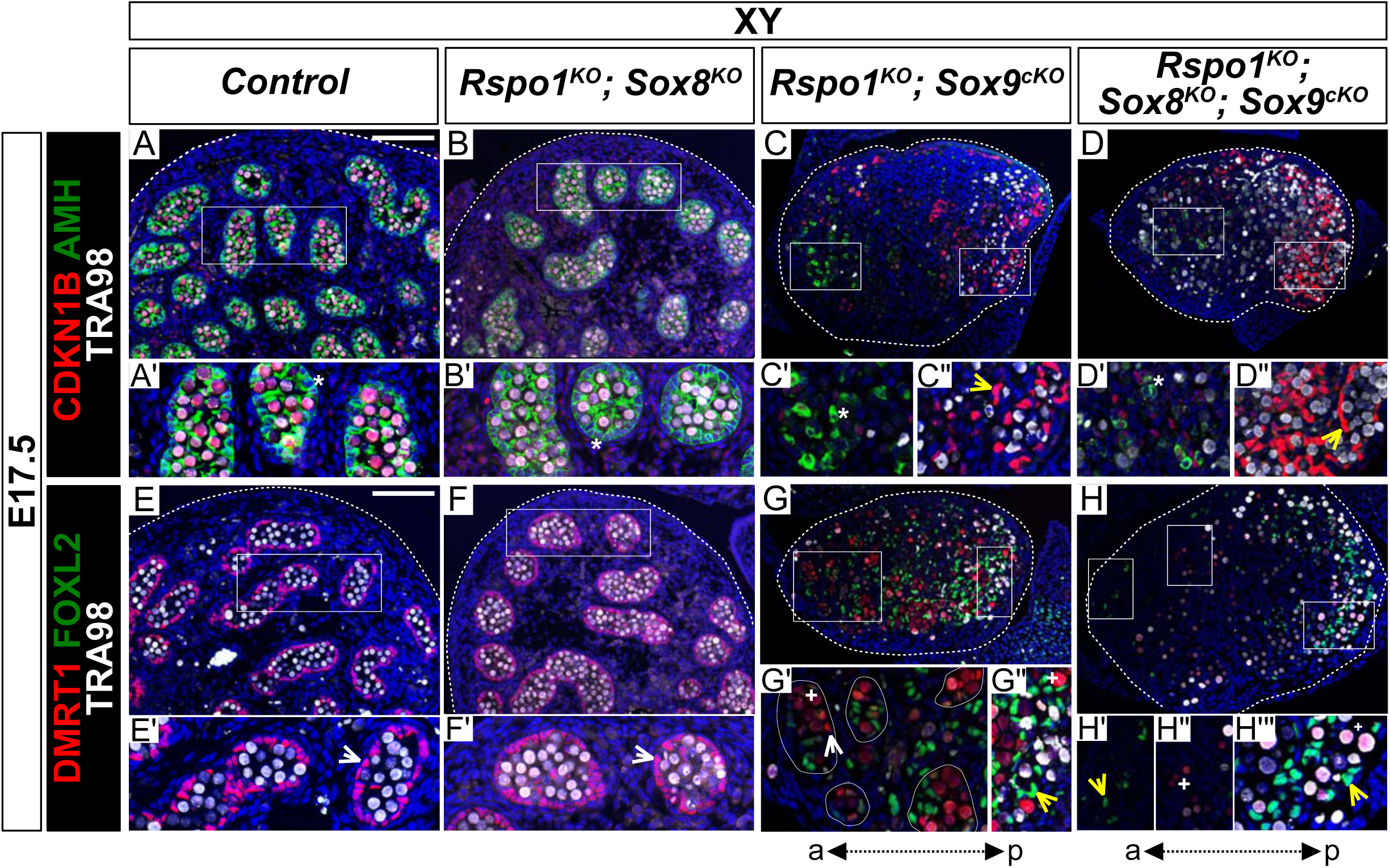
Lack of testis cords in XY *Rspo1^KO^ Sox8^KO^ Sox9^cKO^* triple mutant fetuses at E17.5. Immunofluorescence of CDKN1B (P27) (mitotic arrest marker, in red) (A-D), AMH (Sertoli marker and mature granulosa cell marker, in green) (A-D), DMRT1 (Sertoli and germ cell marker, in red) (E-H), FOXL2 (granulosa cell marker, in green) (E-H), TRA98 (germ cell marker, in white) (A-H), and DAPI (nuclear marker, in blue) (A-H) on gonadal sections from E17.5 fetuses (main panel scale bar 100μm). For gonads in panels C-D, the anterior “a” and posterior “p” axis is shown below each column. Below each main panels A-H, highlighted areas are shown in respective primed letter panels. Yellow arrowheads indicate granulosa cells expressing CDKN1B or FOXL2, asterisks indicate cells expressing AMH, white arrowheads indicate Sertoli cells expressing DMRT1, and plus symbols indicate germ cells expressing DMRT1 and TRA98. Gonads in XY *Rspo1^−/−^*; *Sox8^−/−^* (*Rspo1^KO^ Sox8^KO^*) fetuses exhibited AMH-and DMRT1-positive Sertoli cells organized as testis cords (B, F) and lacked FOXL2-positive granulosa cells (F), as in control testes (A, E). Cells expressing AMH were also were also found in XY *Rspo1^−/−^*; *Sox9^flox/flox^*; *Sf1:cre^Tg/+^* (*Rspo1^KO^ Sox9^cKO^*) and XY *Rspo1^KO^ Sox8^KO^ Sox9^cKO^* gonads (C’ and D’, asterisks), indicating Sertoli cells or mature granulosa cells. Indeed, both exhibited granulosa cells expressing CDKN1B and FOXL2 (C”, D”, G”, H’, H’’’, yellow arrowheads). However, while XY *Rspo1^KO^ Sox9^cKO^* gonads exhibited DRMT1-postive, TRA98-negative Sertoli cells (G’, white arrowhead), these cells were scarce in XY triple mutant gonads (H) (6 of 6 XY triple mutant gonads studied from n=3 fetuses). Note that some DMRT1 expressing cells in XY *Rspo1^KO^ Sox9^cKO^* and XY triple mutant gonads are germ cells expressing TRA98 (G’, G”, H”, H’’’, plus symbols). Thus, although XY *Rspo1^KO^ Sox9^cKO^* and XY triple mutant gonads contain mature granulosa cells at E17.5, these cells do not reprogram as Sertoli cells in XY triple mutant fetuses.

Altogether, fetal XY and XX *Rspo1^KO^ Sox8^KO^ Sox9^cKO^* gonads resembled gonads from XX *Rspo1^KO^ Sox8^KO^*, as well as XY and XX *Rspo1^KO^ Sox9^cKO^* fetuses, with respect to the presence of steroidogenic cells and mature granulosa cells. However, fetal triple mutant gonads lacked Sertoli cells that were present in fetal (*Figure 5F,G*) or post-natal (*Figure 2M,R and Figure 6W*) double mutant mice. Thus, while pre-granulosa cells in triple mutants differentiated precociously, their reprogramming as Sertoli cells forming testis cords at E17.5 appears to be blocked, or delayed.

**Figure 6.**
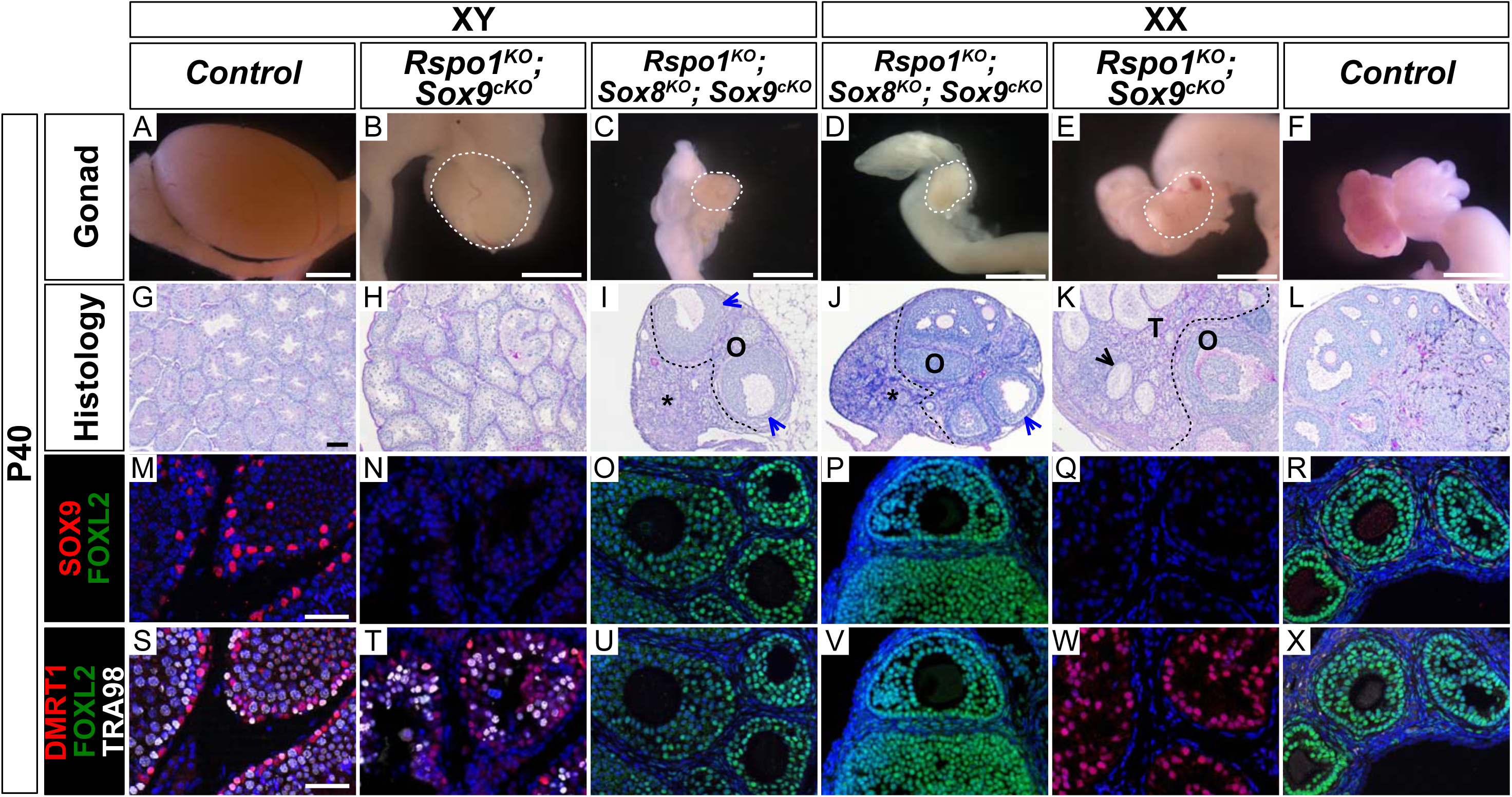
Absence of seminiferous tubules in XY and XX *Rspo1^KO^ Sox8^KO^ Sox9^cKO^* triple mutant adult mice at P40. Macroscopic view of gonads from adult P40 mice (A-F) (Scale bars 1.5mm), histology as revealed by PAS staining on gonadal sections (G-L) (Scale bars 100μm), and immunostainings of SOX9 (Sertoli cell marker, in red) (M-R), FOXL2 (granulosa cell marker, in green) (M-X), DMRT1 (Sertoli and germ cell marker, in red) (S-X), TRA98 (germ cell marker, in white) (S-X), and DAPI (nuclear marker, in blue) (M-X) on gonadal sections (Scale bars 50μm). As shown, adult XY *Rspo1^−/−^*; *Sox9^fl/fl^*; *Sf1:cre^Tg/+^* (*Rspo1^KO^ Sox9^cKO^*) double mutant gonads developed as hypoplastic testes (compare B with A), and XX double mutant gonads developed as ovotestes (E), as in XX *Rspo1^KO^* single mutants and XX *Rspo1^KO^ Sox8^KO^* double mutants (see Figure 2D and 2E). Although *Sox9^fl/fl^* is inactivated by *Sf1:cre^Tg/+^* in *Rspo1^KO^ Sox9^cKO^* mice (N, Q), XY double mutant gonads exhibited seminiferous tubules (H) containing DMRT1-positive Sertoli cells and TRA98-postive germ cells (T), as in control testes (G, S). XX double mutant gonads contained an ovarian compartment “O” with follicles and a testicular “T” compartment with seminiferous tubule-like structures, as indicated by black arrowheads (K). The seminiferous tubule-like structures contained DMRT1-positive Sertoli cells (W), as in control testes (S), but lacked TRA98-postive germ cells (W). In XY and XX *Rspo1^KO^ Sox8^KO^ Sox9^cKO^* triple mutant mice, gonads (C, D) developed as atrophied ovaries (I, J), which were smaller than control ovaries (F, L). XY and XX triple mutant gonads exhibited an ovarian “O” compartment and a distinct interstitial compartment, as indicated by asterisks (I, J). The gonads contained follicles up to the antral stage, though some exhibited irregular granulosa cell organization, as indicated by blue arrowheads (I, J). Notably, XY and XX triple mutants lacked testicular sex cords (I, J) that were present in XY and XX *Rspo1^KO^ Sox9^cKO^* gonads (H, K). Immunostainings on XY and XX *Rspo1^KO^ Sox8^KO^ Sox9^cKO^* gonads confirmed the absence of SOX9-or DMRT1-positive Sertoli cells (O, U, P, V), and the presence of ovarian follicles with granulosa cells expressing FOXL2 (O, U, P, V), as in control ovaries (R, X).

### Lack of Sertoli cell differentiation in XX and XY *Rspo1^KO^ Sox8^KO^ Sox9^cKO^* fetuses

In order to further address the development of triple mutant gonads, we extended our analyses to juvenile (P10) and adult (P40) mice. Both XY and XX *Rspo1^KO^ Sox8^KO^ Sox9^cKO^* triple mutant mice developed externally as female with a short anogenital distance, as in XX control mice *(Figure 6–figure supplement 1C,D,F*). Internally, both XY and XX triple mutants displayed hermaphroditism of the reproductive tracts, as shown by concomitant presence of vasa deferensia and uteri (*Figure 6–figure supplement 1I,J*). This was also observed in XY and XX *Rspo1^KO^ Sox9^cKO^* mice *(Figure 6–figure supplement 1H,K*), as well as in XX *Rspo1^KO^* and in XX *Rspo1^KO^ Sox8^KO^* mice (*Figure 2–figure supplement 1H,I*). Histological analyses revealed that XY and XX triple mutant gonads developed as ovaries containing primary follicles at P10 (*Figure 6–figure supplement 1O,P*), which matured up to the antral follicle stage at P40, though some exhibited irregular granulosa cell organization (*Figure 6I,J, blue arrowheads*). The triple mutant gonads occasionally contained immature or atrophied follicles (*Figure 6–figure supplement 2A,B*). Both XY and XX *Rspo1^KO^ Sox8^KO^ Sox9^cKO^* gonads lacked testicular sex cords (*Figure 6–figure supplement 1O,P; Figure 6I,J; and Figure 6–figure supplement 2A,B*), which were found in XY and XX *Rspo1^KO^* mice lacking *Sox8* (*Figure 2–figure supplement 1L,M and Figure 2G,H*) or *Sox9* (*Figure 6–figure supplement 1N,Q and Figure 6H,K*). Immunostaining experiments on P10 and P40 triple mutant gonads confirmed the presence of follicles with granulosa cells expressing FOXL2 and the absence of testis cords with Sertoli cells expressing SOX9 or DMRT1 (*Figure 6–figure supplement 1U,A’,V,B’ and Figure 6O,U,P,V*).

In 3 of 10 XY and 6 of 16 XX post-natal gonads studied, a cluster of cells expressing DMRT1 were found, but further analyses revealed that these cells did not express the mature Sertoli cell marker GATA1 (Beau, Rauch et al. 2000) (*Figure 6–figure supplement 3F,F’,F’’,L,L’,L’’,O,O’,O’’*). Instead, these cells expressed the embryonic supporting cell marker GATA4 (Tevosian, Albrecht et al. 2002), which suggests rudimentary testis cord formation (*Figure 6–figure supplement 3C,C’,C’’,I,I’,I’’, asterisks*). We also noticed some cells expressing DMRT1 and FOXL2, though these cells were rare (*Figure 6–figure supplement 3L’’,L’’’ arrowheads*). In fact, immunostainings for FOXL2 confirmed that the vast majority of the supporting cells in triple mutants were granulosa cells, which did not undergone reprogramming into Sertoli cells (*Figure 6– figure supplement 1U,A’,V,B’ and Figure 6O,U,P,V,*).

While observing atrophied follicles in adult XY and XX *Rspo1^KO^ Sox8^KO^ Sox9^cKO^* triple mutant mice, a distinct interstitial compartment was also apparent *(Figure 6I,J, asterisks and Figure 6–figure supplement 2A,B*). The identity of this compartment was confirmed by immunostainings for NR5A1 and HSD3β (*Figure 6–figure supplement 2C,D*). In triple mutant gonads, the interstitial cells were arranged individually or in small clusters when compared with XX control ovaries and XX *Rspo1^KO^* ovotestes. In addition, XY and XX triple mutant interstitial cells mildly atrophied, appeared collapsed/dysplastic, and lacked interstitial sinusoids (*Figure 6–figure supplement 2A,B*). No evidence of neoplasia was present in XY and XX triple mutant and in XX *Rspo1^KO^* gonads.

In summary, gonads in XY and XX triple mutants developed as atrophied ovaries. Altogether, our data clearly demonstrate that *Sox8* or *Sox9* is required and sufficient for testicular differentiation in XY and XX *Rspo1^KO^ Sox9^cKO^* or *Rspo1^KO^ Sox8^KO^* double mutants, respectively.

## Discussion

Our results emphasize the essential role of SOX genes in testis differentiation as we show that *Sox* genes are required for Sertoli cell differentiation in XX ovotestis. The critical domain of SOX proteins is the DNA binding domain, the HMG (High-Mobility Group)-domain that binds in a sequence-specific manner (Mertin, McDowall et al. 1999). Remarkably, a male-specific HMG-box gene has been identified in the brown algae *Ectocarpus* (Ahmed, Cock et al. 2014), suggesting a conserved function of the HMG-domain containing genes in maleness throughout evolution. In mice, when the HMG box of SRY is replaced with that of SOX3 or SOX9, these composite *Sox* transgenes induce *Sox9* expression and Sertoli cell differentiation (Bergstrom, Young et al. 2000). Also, transgenic expression of *Sox3* or *Sox10* in XX gonads results in *Sox9* expression and testicular differentiation (Polanco, Wilhelm et al. 2010, Sutton, Hughes et al. 2011). These examples demonstrated functional conservation among *Sox* genes or HMG-box domains and also suggests that male fate centers on transactivation of *Sox9*. However, testicular differentiation was reported in XY and XX *Rspo1*/*Ctnnb1 Sox9* double mutant mice (Lavery, Chassot et al. 2012, Nicol and Yao 2015), suggesting that another *Sox* gene can substitute for the absence of *Sox9* in this context. Given that *Sox8* is up-regulated in the double mutant gonads (Lavery, Chassot et al. 2012, Nicol and Yao 2015), we hypothesized that *Sox8* and *Sox9* can act redundantly for testicular development in mice lacking *Rspo1*. Here, we demonstrated this by showing that in XY and XX *Rspo1^KO^* mice: (*i*) *Sox8* and *Sox9* are expressed independently; (*ii*) *Sox8* or *Sox9* is sufficient for Sertoli cell differentiation in *Rspo1^KO^ Sox9^cKO^* and *Rspo1^KO^ Sox8^KO^* mice, respectively; and (*iii*) *Sox8* and *Sox9* are required for testicular differentiation, as evidenced by the development of atrophied ovaries in *Rspo1^KO^ Sox8^KO^ Sox9^cKO^* triple mutant mice. Together our data show that *Sox8* is able to substitute for *Sox9* to induce Sertoli cell differentiation in XX sex reversal.

The gonad fate in wildtype, *Sox* and *Rspo1* mutant mice is summarized in Figure 6–figure supplement 4. In wildtype mice, SOX9 promotes testicular differentiation in XY gonads and RSPO1 promotes ovarian differentiation in XX gonads (*Figure 6–figure supplement 4A*). This is also the case in mice lacking *Sox8*, since it is dispensable for testis and ovarian development (*Figure 6–figure supplement 4A*) (Sock, Schmidt et al. 2001). As shown, there is an antagonistic relationship between the testis and ovarian pathways, such that the activation of one pathway also leads to the repression of the other to ensure one gonadal fate (*Figure 6–figure supplement 4A*). In XY *Sox9^cKO^* mice, the testis pathway is not activated, and the ovarian pathway is not repressed, leading to ovarian differentiation (*Figure 6–figure supplement 4B*). In XX *Sox9^cKO^* mice, loss of SOX9 does not impair ovarian development (*Figure 6–figure supplement 4B*). In XY *Rspo1^KO^ Sox8^KO^* or XY *Rspo1^KO^ Sox9^cKO^* mice, gonads develop as testes or hypo-plastic testes, since one SOX factor is sufficient for Sertoli cell differentiation and seminiferous tubule formation (*Figure 6– figure supplement 4C,D*). This is also exemplified by ovo-testicular development in XX *Rspo1^KO^ Sox8^KO^* and XX *Rspo1^KO^ Sox9^cKO^* mice (*Figure 6–figure supplement 4C,D*), where Sertoli cells arise from reprogramming of pre-granulosa cells that have precociously differentiated (Maatouk, Mork et al. 2013). We found that inactivation of both SOX factors in mice lacking RSPO1 prevents testicular development in XY and XX animals. In XY and XX *Rspo1^KO^ Sox8^KO^ Sox9^cKO^* triple mutant embryos, though pre-granulosa cells differentiate precociously, the absence of both SOX factors impedes granulosa-to-Sertoli reprogramming in embryos and gonads develop as atrophied ovaries (*Figure 6–figure supplement 4E*). This atrophied ovary outcome suggests that FOXL2 and other ovarian factors cannot fully compensate for the loss of RSPO1 (*Figure 6–figure supplement 4E*).

How *Sox8* operates in pathophysiological cases of testicular differentiation is not yet known. In wildtype mice, *Sox8* expression, like *Sox9*, in XY gonads begins around E11.5 (Jameson, Natarajan et al. 2012, Stevant, Neirijnck et al. 2018). This suggests that SRY may also activate *Sox8*, as predicted (Li, Zheng et al. 2014), or that *Sox8* expression depends on *Sox9*. However, in XY mice lacking the *Sox9* enhancer TESCO, the expression of *Sox9* is reduced in Sertoli cells, but not *Sox8*, indicating that *Sox8* expression does not depend on *Sox9* (Gonen, Quinn et al. 2017). Thus, it is more plausible that SRY activates *Sox8* expression in XY *Rspo1/Ctnnb1 Sox9* double mutant mice. Indeed, in these mice, *Sry* expression is extended beyond E12.5 (Lavery, Chassot et al. 2012, Nicol and Yao 2015), a time when *Sry* is normally down-regulated in mice (Hacker, Capel et al. 1995).

Whereas *Sox8* expression can result from SRY activation in XY embryos, it is not obvious how *Sox8* is upregulated in the absence of *Sry* in XX *Rspo1/Ctnnb1 Sox9* double mutants. One possibility is the involvement of hormons in *Sox8* up-regulation. When E13.5 ovaries are transplanted to kidneys of XY mice, circulating androgens promote partial transdifferentiation towards a testis fate through a mechanism involving up-regulation of *Sox8* before *Sox9* (Miura, Harikae et al. 2019). Ablation of *Sox8* in the transplanted ovaries did not prevent sex reversal and this outcome is likely attributed to the presence of *Sox9*. Furthermore, before up-regulation of *Sox8*, supporting cells in the fetal ovary transplant express *Amh*, a phenotype that is strikingly similar to sex reversal in XX *Rspo1^KO^* gonads (Maatouk, Mork et al. 2013). However, inactivation of *Amh* in the transplanted ovaries was also dispensable for sex reversal, suggesting that TGF-β signaling driven by other TGF-β factors like Activin or unknown factors may promote *Sox* gene expression in sex reversal conditions. The factors controlling *Sox* gene expression in XX transplanted ovaries and in XX mice lacking *Rspo1*/*Ctnnb1* remain to be identified.

The identification of *Sox8* as a key factor in pathophysiological testicular development is somewhat of a paradox, given evidence indicating that aside from *Sry* and *Sox9*, no other *Sox* gene tested so far play key roles in Sertoli cell differentiation in XY wildtype gonads (She and Yang 2017). In mice, *Sox8* is dispensable for Sertoli cell differentiation (Sock Schmidt et al. 2001), but is required for Sertoli cell maintenance along with *Sox9*, since Sertoli cells in XY *Sox8 Sox9* double loss-of-function gonads undergo apoptosis (Barrionuevo, Georg et al. 2009, Barrionuevo, Hurtado et al. 2016). Thus, XY in mice, *Sox8* is genetically redundant for *Sox9* during sex determination, after which both *Sox* play synergistic roles in Sertoli cells. However, functional redundancy between SOX8 and SOX9 does not seem to operate in humans given that mutations of SOX8 were associated with a range of phenotypes including complete gonadal dysgenesis (streak gonads with immature female genitalia) and hypoplastic testes in three 46, XY patients (Portnoi, Dumargne et al. 2018). Thus, in humans, SOX8 is emerging to be an important regulator of testicular gonadal development and by extension, overall male development. In mice, our data here demonstrates that *Sox8* plays an essential role for testicular differentiation in sex reversal conditions.

## Materials and Methods

### Mouse strains and genotyping

The experiments described here were carried out in compliance with the relevant institutional and French animal welfare laws, guidelines, and policies. These procedures were approved by the French ethics committee (Comité Institutionnel d’Ethique Pour l’Animal de Laboratoire; number NCE/2011-12). All mouse lines were kept on a mixed 129Sv/C57BL6/J background. *Rspo1^−/−^* (Chassot, Ranc et al. 2008), *Sox8^−/−^* (Sock, Schmidt et al. 2001), *Sox9^fl/fl^* (Akiyama, Chaboissier et al. 2002), and *Sf1:cre^Tg/+^* (Bingham, Verma-Kurvari et al. 2006) mice were obtained previously, and the generation of *Sox9^fl/fl^*; *Sf1:cre^Tg/+^* (Lavery, Lardenois et al. 2011) and *Rspo1^−/−^*; *Sox9^fl/fl^*; *Sf1:cre^Tg/+^* (Lavery, Chassot et al. 2012) mice was described previously. For *Rspo1^KO^ Sox8^KO^* mice: *Rspo1^−/−^* males were mated with *Sox8^−/−^* females to obtain *Rspo1^+/−^*; *Sox8^+/−^* males and females. Matings between these littermates allowed us to obtain *Rspo1^−/−^*; *Sox8^−/−^* double mutant mice, referred to as *Rspo1^KO^ Sox8^KO^* mice, and control animals. For *Rspo1^KO^ Sox8^KO^ Sox9^cKO^* mice: first, *Rspo1^−/−^*; *Sox8^−/−^* males were mated with *Sox8^−/−^*; *Sox9^fl/fl^*; *Sf1:cre^Tg/+^* females to generate *Rspo1^+/−^*; *Sox8^−/−^*; *Sox9^fl/+^* males and *Rspo1^+/−^*; *Sox8^−/−^*; *Sox9^fl/+^*; *Sf1:cre^Tg/+^* females. Matings between these littermates then produced *Rspo1^−/−^*; *Sox8^−/−^*; *Sox9^fl/fl^* males and *Rspo1^+/−^*; *Sox8^−/−^*; *Sox9^fl/fl^*; *Sf1:cre^Tg/+^* females. Finally, matings between these littermates then produced allowed us to obtain *Rspo1^−/−^*; *Sox8^−/−^*; *Sox9^fl/fl^*; *Sf1:cre^Tg/+^* triple mutant mice, referred to as *Rspo1^KO^ Sox8^KO^ Sox9^cKO^* mice, and control animals. Embryos were collected from timed evening matings that was confirmed by the presence of a vaginal plug the following morning. This marked embryonic day 0.5 (E0.5). The day of delivery was defined as post-natal day 0 (P0). Genotyping was performed as described in (Chaboissier, Kobayashi et al. 2004, Bingham, Verma-Kurvari et al. 2006, Chassot, Ranc et al. 2008) by using DNA extracted from tail tip or ear biopsies of mice. The presence of the Y chromosome was determined, as described previously (Hogan 1994).

### *In situ* hybridization

Gonad samples were fixed with 4% paraformaldehyde overnight, processed for paraffin embedding, and then sectioned at 5-7 μm thick. The *in situ* hybridizations for Figure 1E-H were carried out essentially as described by (Lavery, Chassot et al. 2012). For analyses in Figure 1A-D and Figure 1–figure supplement 1A, RNAscope technology was used (Wang, Flanagan et al. 2012). The *Rspo1* probe was purchased from the manufacturer (Advanced Cell Diagnostics) and the protocol was performed according to the manufacturer’s instructions using the Fast Red dye, which can be visualized using light or fluorescence microscopy. The *in situ* hybridization experiments were performed on gonads from at least three mice for each genotype.

### X-gal staining

Gonad samples were fixed with 4% paraformaldehyde for up to 4 hours, processed for frozen embedding, and then cryosectioned at 10 μm thick. Stainings were carried out essentially as described by (Chassot, Gregoire et al. 2011). Xgal stainings were performed on gonads from at least three mice for each genotype.

### Immunological analyses

Gonad samples were fixed with 4% paraformaldehyde overnight, processed for paraffin embedding, and sectioned at 5 μm thick. The following dilutions of primary antibodies were used: AMH/MIS (c-20, sc-6886, Santa Cruz), 1:200; DMRT1 (HPA027850, Sigma), 1:100; FOXL2 (NB100-1277, Novus), 1:200; GATA1 (N6, sc-265, Santa Cruz), 1:200; GATA4 (C20, sc-1237, Santa Cruz), 1:200; 3βHSD (P18, sc-30820, Santa Cruz), 1:200; P27 (Kip1, sc-528, Santa Cruz), 1:200; SF1 (kindly provided by Ken Morohashi), 1:1000; SOX9 (HPA001758, Sigma), 1:200; and TRA98 (ab82527, Abcam), 1:200. Counterstain with DAPI was used to detect nuclei. Immunofluorscence of secondary antibodies were detected with an Axio ImagerZ1 microscope (Zeiss) coupled to an Axiocam mrm camera (Zeiss). Images were processed with Axiovision LE and Serif Affinity Photo software. Immunostaining experiments were performed on gonads from at least three mice for each genotype.

### Cell quantification

Immunostaining analyses were performed, as described above. For analyses at E17.5, immunostainings were performed on 2 to 17 sections spaced 20-30μm apart in each gonad. Then, for each section, the ratio of cells positive for DMRT1, FOXL2, or CDKN1B to total gonad area, as visualized by DAPI staining, were manually tabulated. Finally, the ratios and mean for each genotype were plotted in a histogram using Graphpad software.

### Histological analyses

Gonad samples were fixed with Bouin’s solution overnight, processed for paraffin embedding, sectioned at 5 μm thick, and then stained according to standard procedures for periodic acid Schiff (PAS) or hematoxylin and eosin (H&E) staining. Images were taken with an Axiocam mrm camera (Zeiss) and processed with Serif Affinity Photo software. Histology staining was performed on gonads from at least three mice for each genotype.

## Acknowledgements

We thank Ken Morohashi for the SF1/NR5A1 antibody. We also thank Samah Rekima and the Experimental Histopathology Platform for assistance with samples. Thanks to the Schedl team for helpful discussions, and Anne-Amandine Chassot and Aitana Perea-Gomez for critical reading of the manuscript. This work was supported by the Agence Nationale de la Recherche (ANR-11-LABX-0028-01; ANR-19-CE14-0022-*SexDiff*).

---

**Figure 1–figure supplement. 1.**
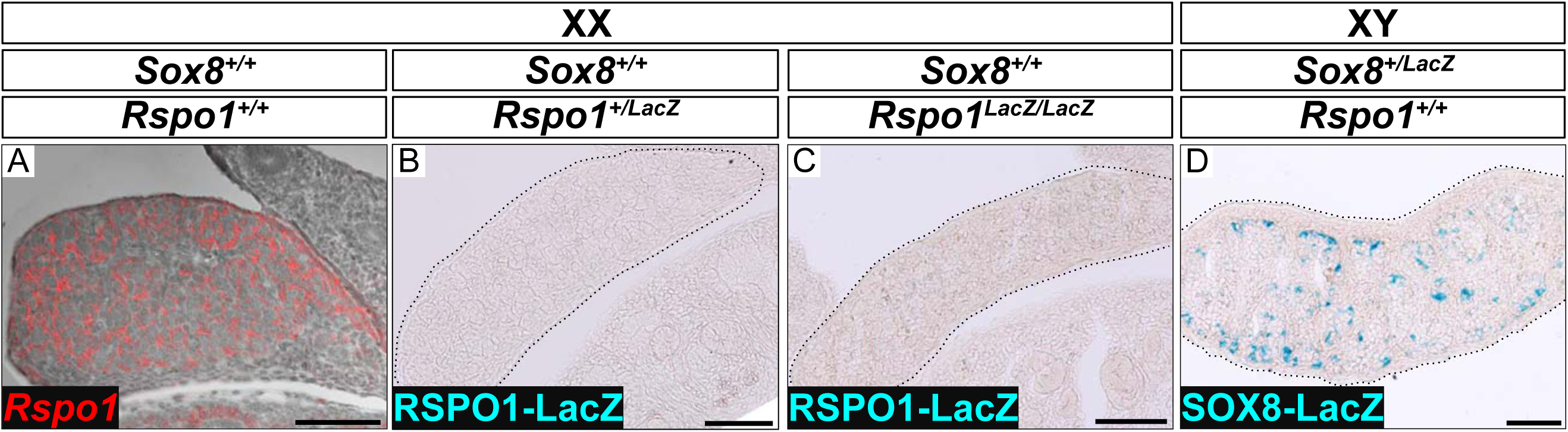
Absence of β-galactosidase activity in fetal XX *Rspo1^+/−^* and *Rspo1^−/−^* gonads. In situ hybridization experiments using a *Rspo1* anti-sense probe on XX wild-type ovaries at E14.5 shows robust *Rspo1* expression (A). In contrast, detection of RSPO1-LacZ by β-galactosidase staining on XX heterozygous *Rspo1^+/−^* (*Rspo1^LacZ/+^*) and homozygous *Rspo1^−/−^* (*Rspo1^LacZ/LacZ^*) E13.5 gonads shows no staining (B, C). This is because the RPSO1-LacZ protein is not retained in cells. Detection of SOX8-LacZ in testes of E13.5 XY *Sox8^+/−^* (*Sox8^LacZ/+^*) littermates (D) is a positive control for the β-galactosidase staining. Scale bars 100μm.

**Figure 2–figure supplement 1.**
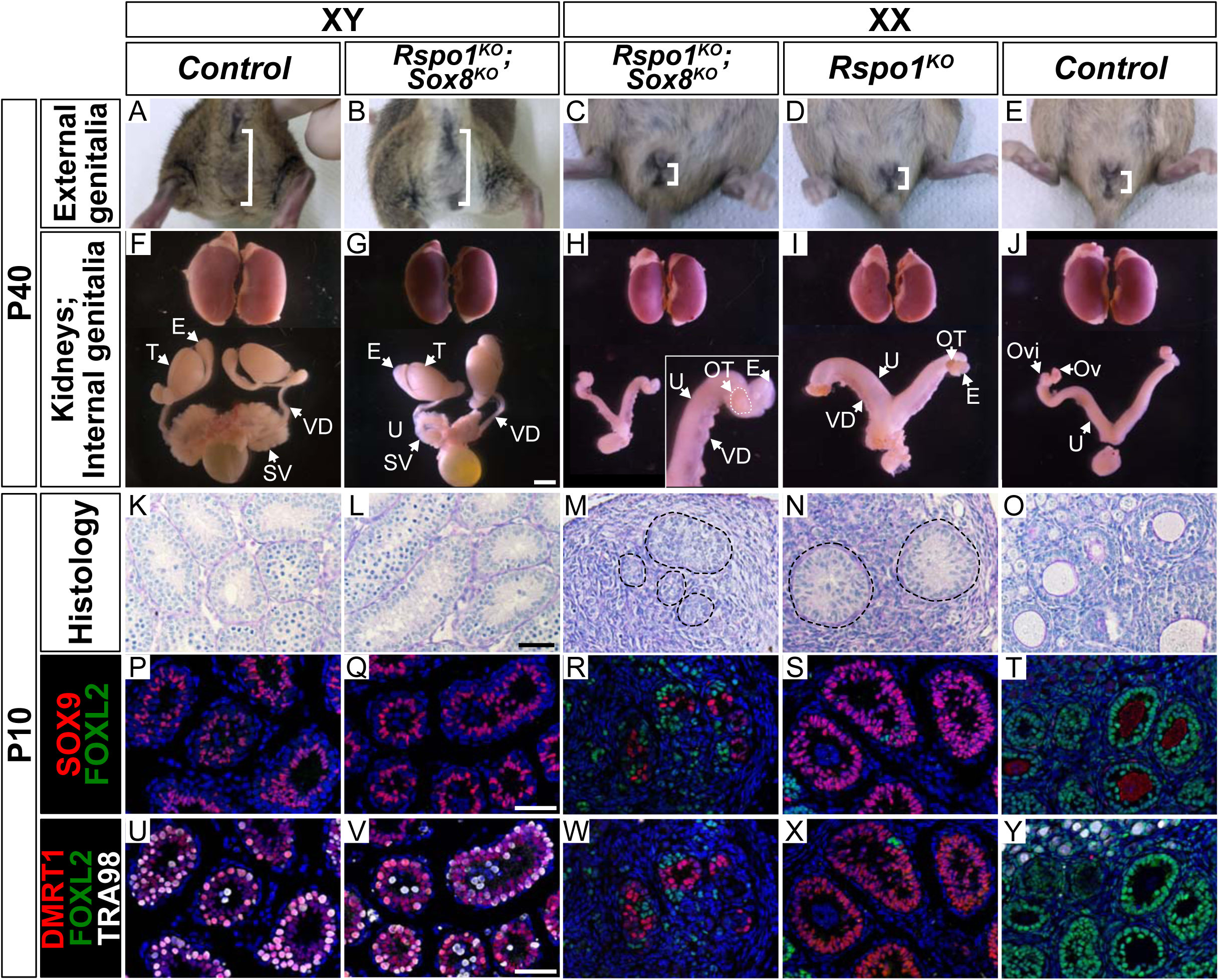
Secondary sex characteristics of XY and XX *Rspo1^KO^ Sox8^KO^* adult P40 mice and analyses in juvenile P10 mice. Externally, at P40, XY *Rspo1^KO^ Sox8^KO^* mice developed a long anogenital distance (B), as in XY control males (A), whereas XX control females and XX *Rspo1^KO^* mice developed a short anogenital distance (D, E). Internally, XY control mice and XY *Rspo1^KO^ Sox8^KO^* mice developed as male with testes “T”, epididymides “E”, vasa deferentia “VD”, and seminal vesicles “SV” (F, G). XX control mice developed as female with ovaries “Ov”, oviducts “Ovi”, and uteri “U” (H), as in control females (J). Both XX *Rspo1^KO^* mice and XX *Rspo1^KO^ Sox8^KO^* mice exhibited hermaphroditism of the reproductive tracts (H, I). Kidneys are shown for comparison (F-J). In P10 mice, XY control and XY *Rspo1^KO^ Sox8^KO^* gonads exhibited seminiferous tubules (K, L) containing SOX9-and DMRT1-positive Sertoli cells (P, Q, U, V). XX control ovaries exhibited follicles (O) containing FOXL2-postive granulosa cells (T, Y). Gonads in XX *Rspo1^KO^* and XX *Rspo1^KO^ Sox8^KO^* mice contained SOX9-and DMRT1-postive Sertoli cells (R, S, W, X), indicating XX sex reversal. As shown in Figure 2, ovotestis development is indistinguishable in XX *Rspo1^KO^* and XX *Rspo1^KO^ Sox8^KO^* mice by P40. Internal organs scale bar 3mm (F-J), histology and immunostainings scale bar 50μm (K-Y).

**Figure 3–figure supplement 1.**
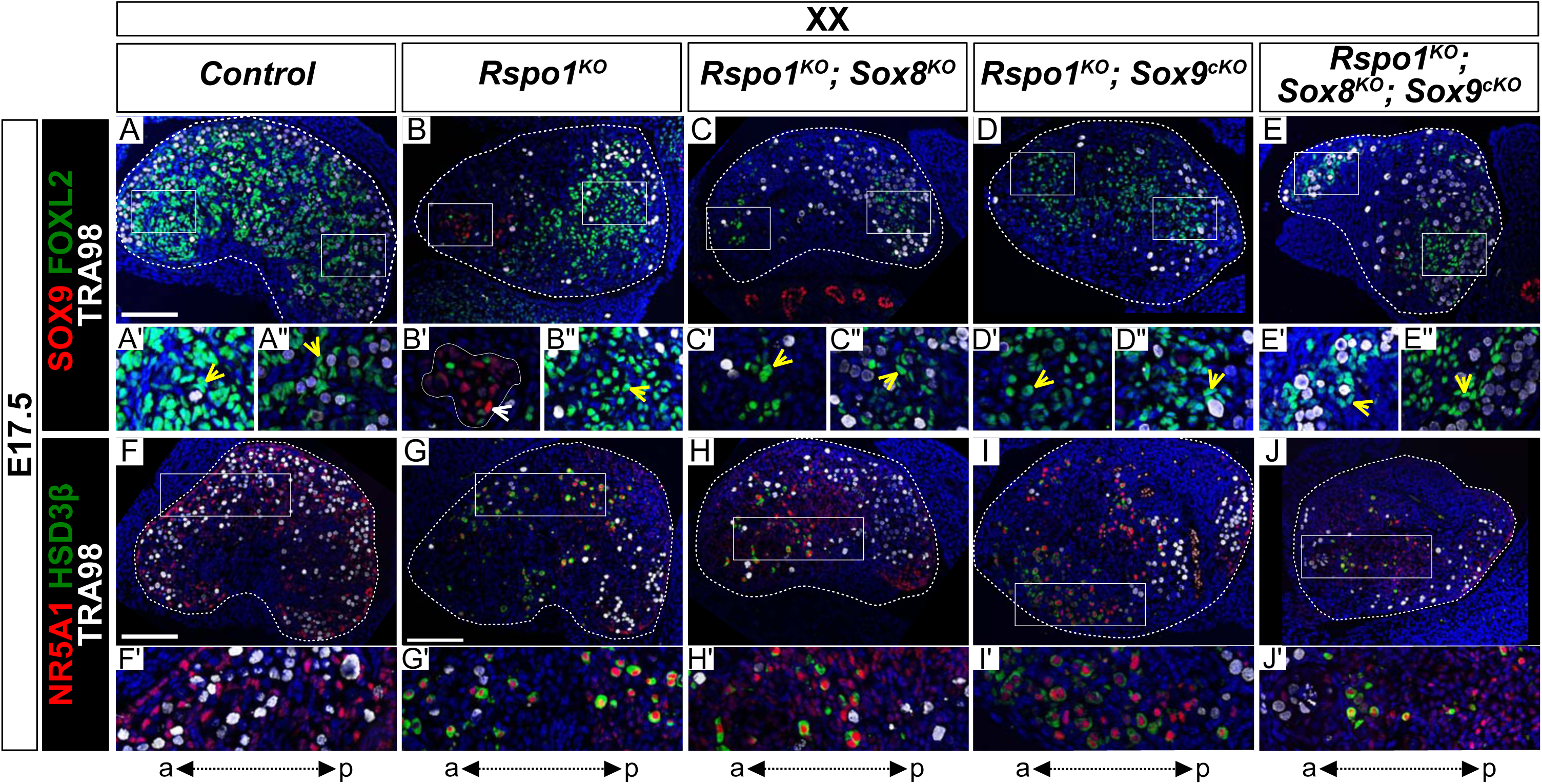
Absence of SOX9 expression in gonads from XX *Rspo1^KO^ Sox8^KO^ Sox9^cKO^* fetuses and presence of steroidogenic cells at E17.5. Immunofluorescence of SOX9 (Sertoli cell marker, in red) (A-E), FOXL2 (granulosa cell marker, in green) (A-E), NR5A1 (SF1) (supporting and steroidogenic cell marker, in red) (F-J), HSD3B1/2 (HSD3β) (F-J), (steroidogenic cell marker, in green), TRA98 (germ cell marker, in white) (A-J), and DAPI (nuclear marker, in blue) (A-T) on gonadal sections from E17.5 fetuses (main panel scale bar 100μm). The anterior “a” and posterior “p” axis is shown below each column. For main panels A-E, highlighted anterior and posterior areas are shown in the respective single and double primed letter panels. Yellow arrowheads indicate granulosa cells expressing FOXL2 and white arrowheads indicate Sertoli cells expressing SOX9. Gonads in XX control fetuses developed as ovaries, as shown by FOXL2 expression in pre-granulosa cells (A’, yellow arrowhead). In contrast, XX *Rspo1^−/−^* (*Rspo1^KO^*) gonads exhibited Sertoli cells expressing SOX9 forming testis cords (B’, white arrowhead). At this stage, XX *Rspo1^−/−^*; *Sox8^−/−^* (*Rspo1^KO^ Sox8^KO^*), XX *Rspo1^−/−^*; *Sox9^flox/flox^*; *Sf1:cre^Tg/+^* (*Rspo1^KO^ Sox9^cKO^*), and XX *Rspo1^KO^ Sox8^KO^ Sox9^cKO^* gonads lacked SOX9-positive Sertoli cells because sex reversal is delayed (C), or because of genetic inactivation *Sox9^flox/flox^* with *Sf1:cre^Tg/+^* (D, E). The XX single, double, and triple mutant gonads contained steroidogenic cells expressing NR5A1 and HSD3β (G’-J’), which were absent in control fetal ovaries (F’).

**Figure 5–figure supplement 1.**
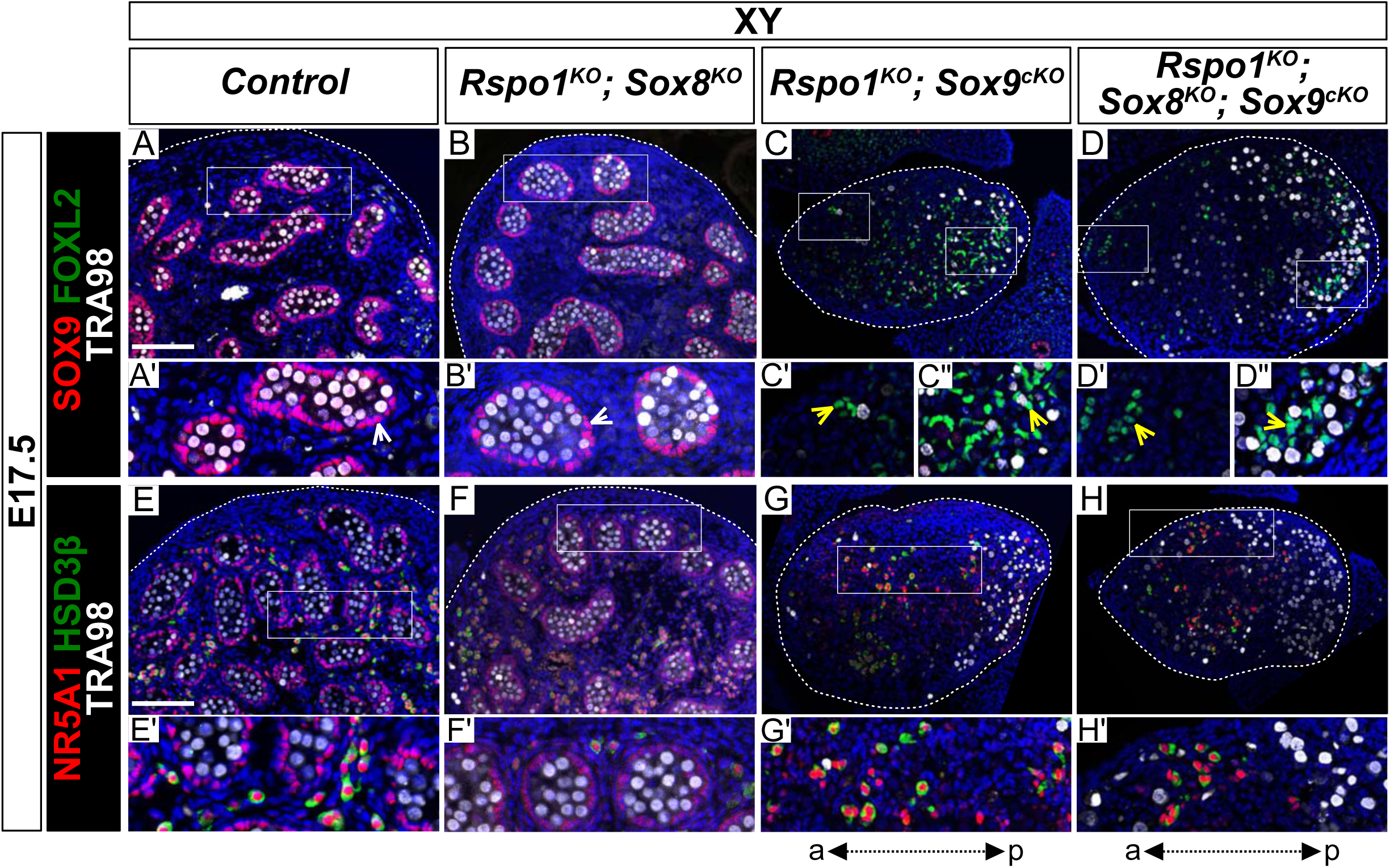
Absence of SOX9 expression in gonads from XY *Rspo1^KO^ Sox8^KO^ Sox9^cKO^* fetuses and presence of steroidogenic cells at E17.5. Immunofluorescence of SOX9 (Sertoli cell marker, in red) (A-D), FOXL2 (granulosa cell marker, in green) (A-D), NR5A1 (SF1) (supporting and steroidogenic cell marker, in red) (E-H), HSD3B1/2 (HSD3β) (E-H), (steroidogenic cell marker, in green), TRA98 (germ cell marker, in white) (A-H), and DAPI (nuclear marker, in blue) (A-H) on gonadal sections from E17.5 fetuses (main panel scale bar 100μm). For gonads in C-D and G-H, the anterior “a” and posterior “p” axis is shown below each column. Also, for main panels in A-H, highlighted areas are shown in the respective single and double primed letter panels. Yellow arrowheads indicate granulosa cells expressing FOXL2 and white arrowheads indicate Sertoli cells expressing SOX9. Gonads in XY *Rspo1^−/−^*; *Sox8^−/−^* (*Rspo1^KO^ Sox8^KO^*) fetuses exhibited SOX9-positive Sertoli cells organized as testis cords and lacked FOXL2-positive granulosa cells (B, B’), as in control testes (A, A’). As in XY *Rspo1^KO^ Sox9^cKO^* gonads (C), genetic inactivation of *Sox9* in XY *Rspo1^KO^ Sox8^KO^ Sox9^cKO^* gonads resulted in the absence of SOX9-positive Sertoli cells (D). However, these gonads exhibited granulosa cells expressing FOXL2 (C” and D”, asterisks). As in fetal testes (E, F), XY *Rspo1^KO^ Sox9^cKO^* and XY triple mutant gonads exhibited steroidogenic cells expressing NR5A1 and HSD3β (G, H).

**Figure 6–figure supplement 1.**
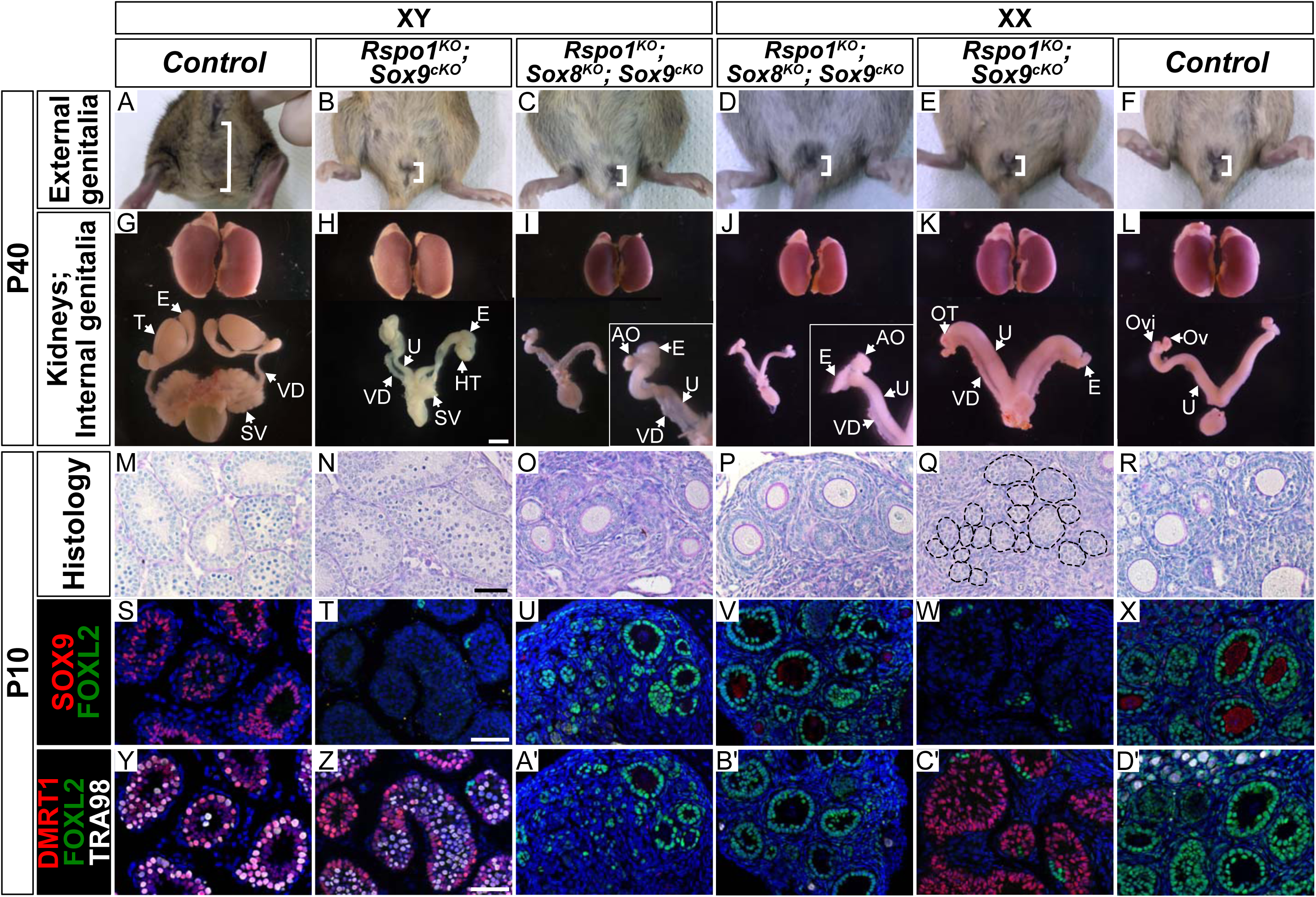
Secondary sex characteristics of XY and XX *Rspo1^KO^ Sox8^KO^ Sox9^cKO^* adult P40 mice and analyses in juvenile P10 mice. Externally, at P40, XY and XX *Rspo1^KO^ Sox8^KO^ Sox9^cKO^* mice (C, D), and XY and XX *Rspo1^KO^ Sox9^cKO^* mice (B, E) developed a short anogenital distance, as in XX control females (F). Internally, XX control females developed ovaries “Ov”, oviducts “Ovi”, and uteri “U” (L). In contrast, XY control mice developed a long anogenital distance (A) and internally contained testes “T”, epididymides “E”, vasa deferentia “VD”, and seminal vesicles “SV” (G). Both XY and XX triple mutants exhibited hermaphroditism of the reproductive tracts (I, J), like XY and XX *Rspo1^KO^ Sox9^cKO^* double mutant mice (H, K). However, while XY *Rspo1^KO^ Sox9^cKO^* mice developed testes “HT” that were hypoplastic (H) and XX *Rspo1^KO^ Sox9^cKO^* mice developed ovotestes “OT” (K), XY and XX triple mutants developed atrophied ovaries “AO” (I, J). Kidneys are shown for comparison (G-L). In P10 mice, XY and XX triple mutants exhibited ovarian follicles (O, P) containing FOXL2-positive granulosa cells (U, A’, V, B’) like XX control ovaries (R, X, D’). This contrasts gonad development in XY and XX *Rspo1^KO^ Sox9^cKO^* gonads, which contained seminiferous tubules (N) or seminiferous tubule-like structures (Q, circled) with DMRT1-positive Sertoli cells (Z, C’), similar to XY control testes (M, S, Y). Internal organs scale bar 3mm (G-L), histology and immunostainings scale bar 50μm (M-D’).

**Figure 6–figure supplement 2.**
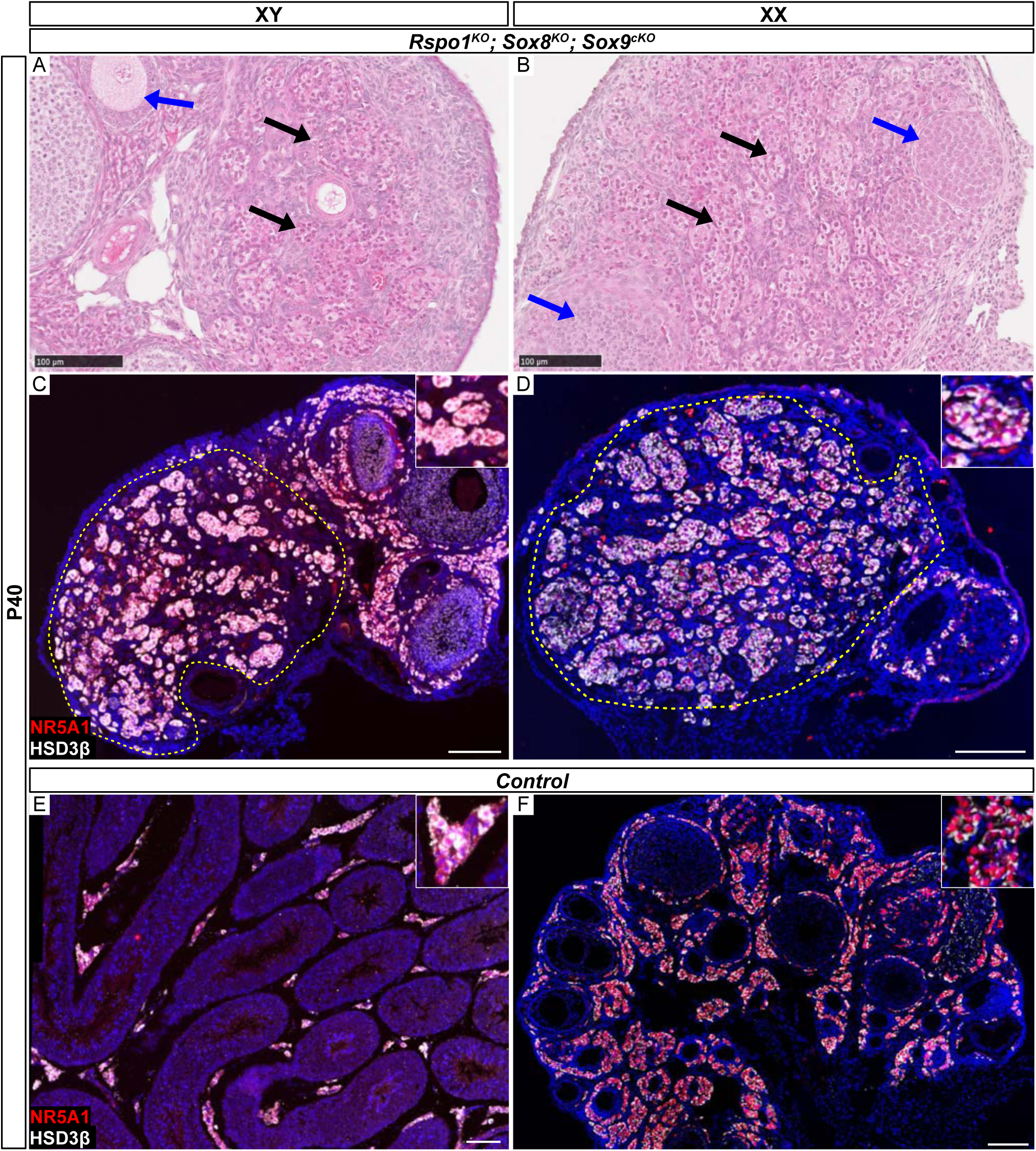
Organization of adult XY and XX *Rspo1^KO^ Sox8^KO^ Sox9^cKO^* triple mutant gonads. Histology (H&E) staining of adult P40 XY and XX *Rspo1^KO^ Sox8^KO^ Sox9^cKO^* triple mutant gonads show collapsed/atrophied interstitial cells (black arrows) and immature/atrophied follicles (blue arrows) (A, B). Immunofluorescence of NR5A1 (somatic cell marker, in red) and HSD3B1/2 (HSD3β) (interstitial/steroidogenic cell marker, in white) reveals a compartment of interstitial cells (circled). In control testes or ovaries, interstitial cells are found between seminiferous tubules or follicles (E, F). All scale bars 100μm.

**Figure 6–figure supplement 3.**
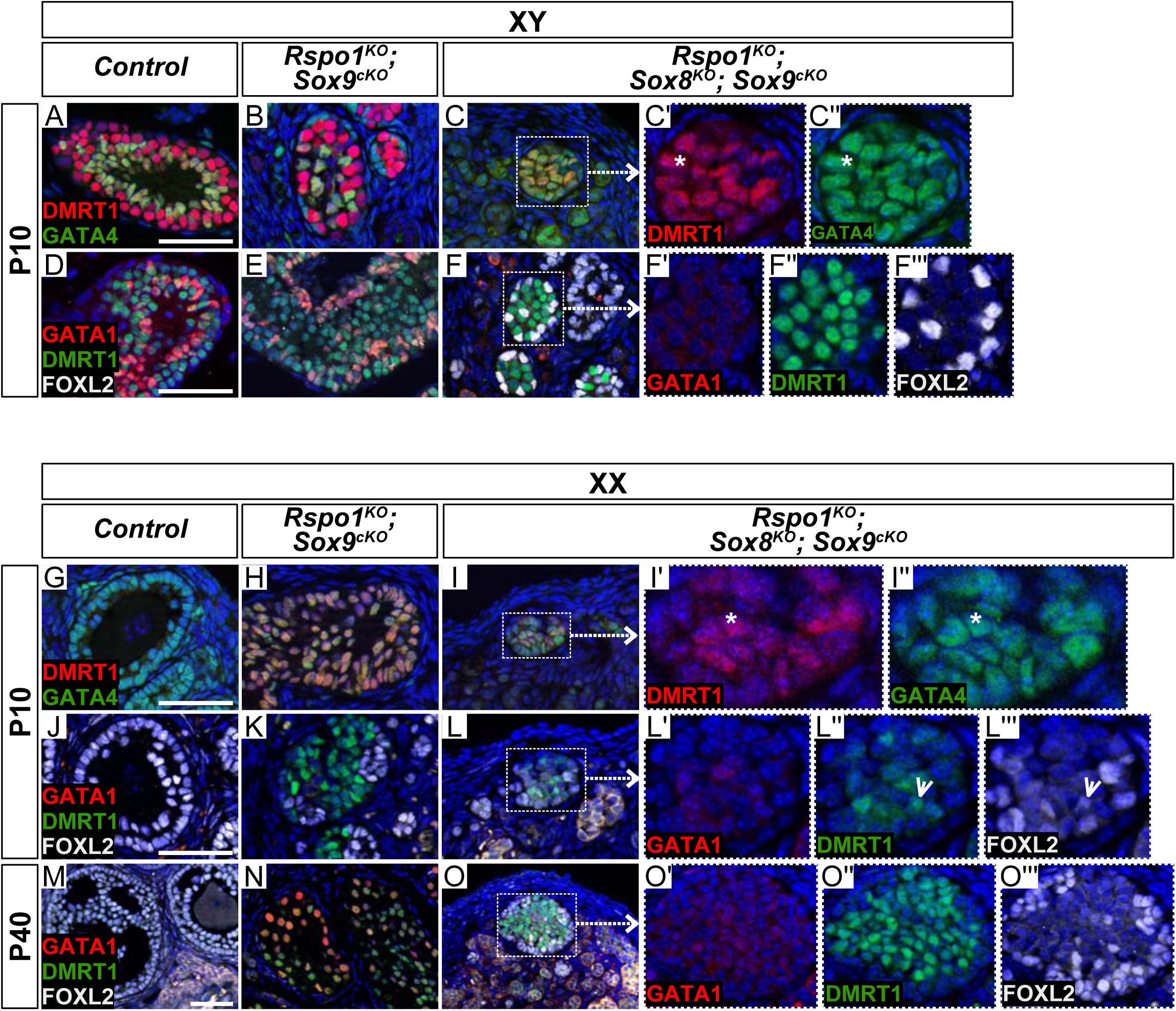
Rare immature Sertoli cells in XY and XX *Rspo1^KO^ Sox8^KO^ Sox9^cKO^* triple mutant gonads. Both XY or XX triple mutants exhibited rare cells expressing DMRT1, a Sertoli cell marker, and GATA4, a somatic cell marker (asterisks in C’ and C’’, and in I’ and I’’), but lacked DMRT1-postive cells Sertoli cells expressing the mature Sertoli cell marker GATA1 (F’, L’, O’). Cells expressing DMRT1 (F, L, O) were sometimes found in close proximity with cells expressing the granulosa cell marker FOXL2 (F’’, F’’’, L’’, L’’’, O’’, O’’’), and some cells co-expressed DMRT1 and FOXL2 (L”, L”’, arrowheads). Though GATA1-postive cells are absent in XX *Rspo1^KO^ Sox9^cKO^* mice at P10 (E), they are present in P40 gonads (K, N). Scale bars 50μm.

**Figure 6–figure supplement 4.**
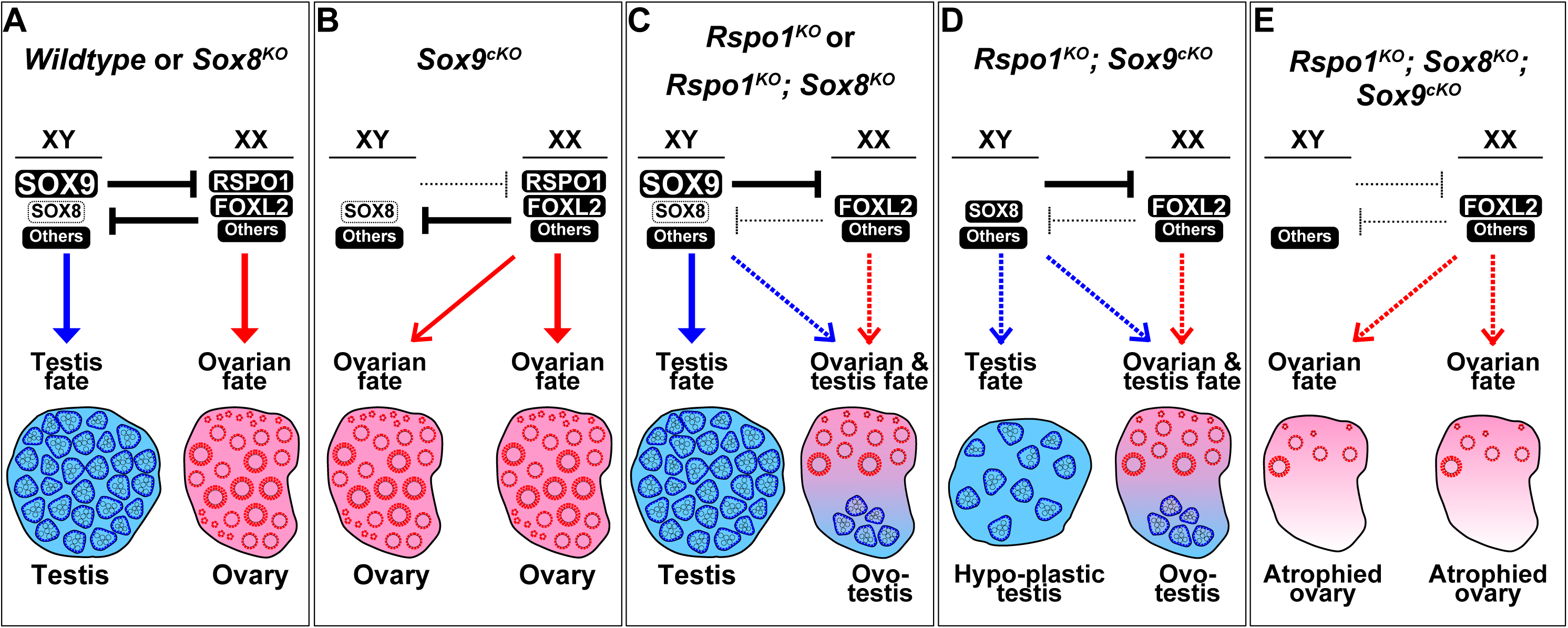
Gonad fate in wildtype, *Rspo1*, and *Sox* mutant mice. In wildtype mice, SOX9, SOX8, and other factors promote testicular differentiation in XY mice, and RSPO1, FOXL2, and other factors promote ovarian differentiation in XX mice, as indicated by arrows (A). Antagonism exists between the testis and ovarian pathway, as indicated by “T” bars (A). SOX9 and RSPO1 are essential for testicular and ovarian differentiation respectively, since XY *Sox9^cKO^* mice develop ovaries (B) and XX *Rspo1^KO^* mice develop partial sex-reversal ovotestes (C). However, we previously demonstrated that SOX9 is dispensable for testicular development in XX *Rspo1^KO^* mice, by studying XX *Rspo1^KO^ Sox9^cKO^* mice (D). Also, gonads in XY *Rspo1^KO^ Sox9^cKO^* mice develop as hypoplastic testes (D), indicating that RSPO1 is required for ovarian differentiation in XY *Sox9^cKO^* mice. In studying a *Sox8^KO^* mutation in XY and XX mice (A) and *Rspo1^KO^* (C) mice, it was clear that SOX8 is dispensable for testicular, ovarian, or ovotesticular development. In this study however, we demonstrated that SOX8 is required for hypoplastic testicular or ovotesticular differentiation in XY and XX *Rspo1^KO^ Sox9^cKO^* mice (D) by studying triple mutants (E). Gonads in both XY and XX *Rspo1^KO^ Sox8^KO^ Sox9^cKO^* mice lacked testis cords and developed as atrophied ovaries (E). Thus, SOX8 or SOX9 is sufficient and both SOX are required for testicular differentiation in gonads lacking RSPO1.

## Notes

#### Summary of Updates

The list of the authors has been updated

